# Highly Specific Enrichment of Rare Nucleic Acids using *Thermus Thermophilus* Argonaute with Applications in Cancer Diagnostics

**DOI:** 10.1101/491738

**Authors:** Jinzhao Song, Jorrit W. Hegge, Michael G. Mauk, Junman Chen, Jacob E. Till, Neha Bhagwat, Lotte T. Azink, Jing Peng, Moen Sen, Jazmine Mays, Erica Carpenter, John van der Oost, Haim H. Bau

**Author notes:** These authors contributed equally to this work (Jinzhao Song, Jorrit W. Hegge).

## Abstract

Detection of disease-associated, cell-free nucleic acids enables early diagnostics, genotyping, and personalized therapy, but is challenged by their low concentration and sequence homology with abundant wild-type nucleic acids. We describe a novel approach, dubbed NAVIGATER, for **<u>N</u>**ucleic **<u>A</u>**cid enrichment **<u>V</u>**ia DNA-**<u>G</u>**uided **<u>A</u>**rgonaute from ***<u>T</u>**hermus th**<u>er</u>**mophilus* (*Tt*Ago) that allows for specific cleavage of guide-complementary DNA and RNA with single nucleotide precision. NAVIGATER greatly increases the fractions of rare alleles, enhancing the sensitivity of downstream detection such as ddPCR, sequencing, and clamped enzymatic amplification. We demonstrated 60-fold enrichment of *KRAS* G12D and nearly 100-fold increased sensitivity of clamped-PCR (PNA and XNA-PCR), enabling detection of low-frequency (0.01%) mutant alleles (∼1 copy) in blood samples of pancreatic cancer patients. NAVIGATER surpasses Cas9-DASH, identifying more mutation-positive samples when combined with XNA-PCR. Moreover, *Tt*Ago does not require the target to contain any specific (PAM-like) motifs; is a multi-turnover enzyme; cleaves ssDNA, dsDNA, and RNA targets in a single assay; and operates at elevated temperatures, providing high selectivity and compatibility with polymerases. The here-described NAVIGATER approach has important advantages over other enrichment methods.

## Introduction

In recent years, researchers have identified various enzymes from archaea and bacteria that can be programmed with nucleic acids to cleave complementary strands, among which CRISPR-Cas have attracted considerable attention as genome-editing tools,^1-3^. In medical diagnostics, CRISPR-associated nucleases have been used to (1) amplify reporter signals during nucleic acid detection^4-8^, (2) detect unamplified targets immobilized on graphene field-effect transistors,^9^, and (3) enrich rare alleles to enhance detection limits of oncogenic sequences^10-12^. Despite remarkable progress, the applications of CRISPR-Cas are restricted due, among other things, to their reliance on the protospacer-adjacent motif (PAM), which is absent in many sequences of interest^10-12^.

Analogous to CRISPR-Cas, Argonaute (Ago) proteins^13^ are nucleic acid-guided endonucleases. In contrast to Cas nucleases, Ago nucleases do not require the presence of any specific motifs and are, therefore, more versatile. We found that under appropriate conditions, Ago from the thermophilic bacterium *Thermus thermophilus* (*Tt*Ago)^14-17^ cleaves with high efficiency both DNA and RNA complimentary to its DNA guide (gDNA), but spares nucleic acids with a single nucleotide mismatch at and around its catalytic site. We formulate a new rare allele enrichment assay (Fig. 1) termed NAVIGATER (**<u>N</u>**ucleic **<u>A</u>**cid enrichment **<u>V</u>**ia DNA **<u>G</u>**uided **<u>A</u>**rgonaute from ***<u>T</u>**hermus th**<u>er</u>**mophilus*). We optimize guide length, assay composition, and incubation temperature to enable NAVIGATER to increase rare allele fractions of *KRAS, EGFR, and BRAF* mutants in blood samples from cancer patients, greatly improving the sensitivity of downstream detection schemes such as droplet digital PCR (ddPCR)^18^, Peptide Nucleic Acid-Mediated PCR (PNA-PCR)^19^, PNA-Loop Mediated Isothermal Amplification (LAMP)^20^, Xenonucleic Acid clamp PCR (XNA-PCR)^21^, and Sanger sequencing. NAVIGATER has the potential to greatly increase the sensitivity and clinical utility of non-invasive tests such as liquid biopsy, especially for monitoring of somatic mutations through cell-free DNA (cfDNA) for early diagnostics and for personalized therapy.

**Fig. 1.**
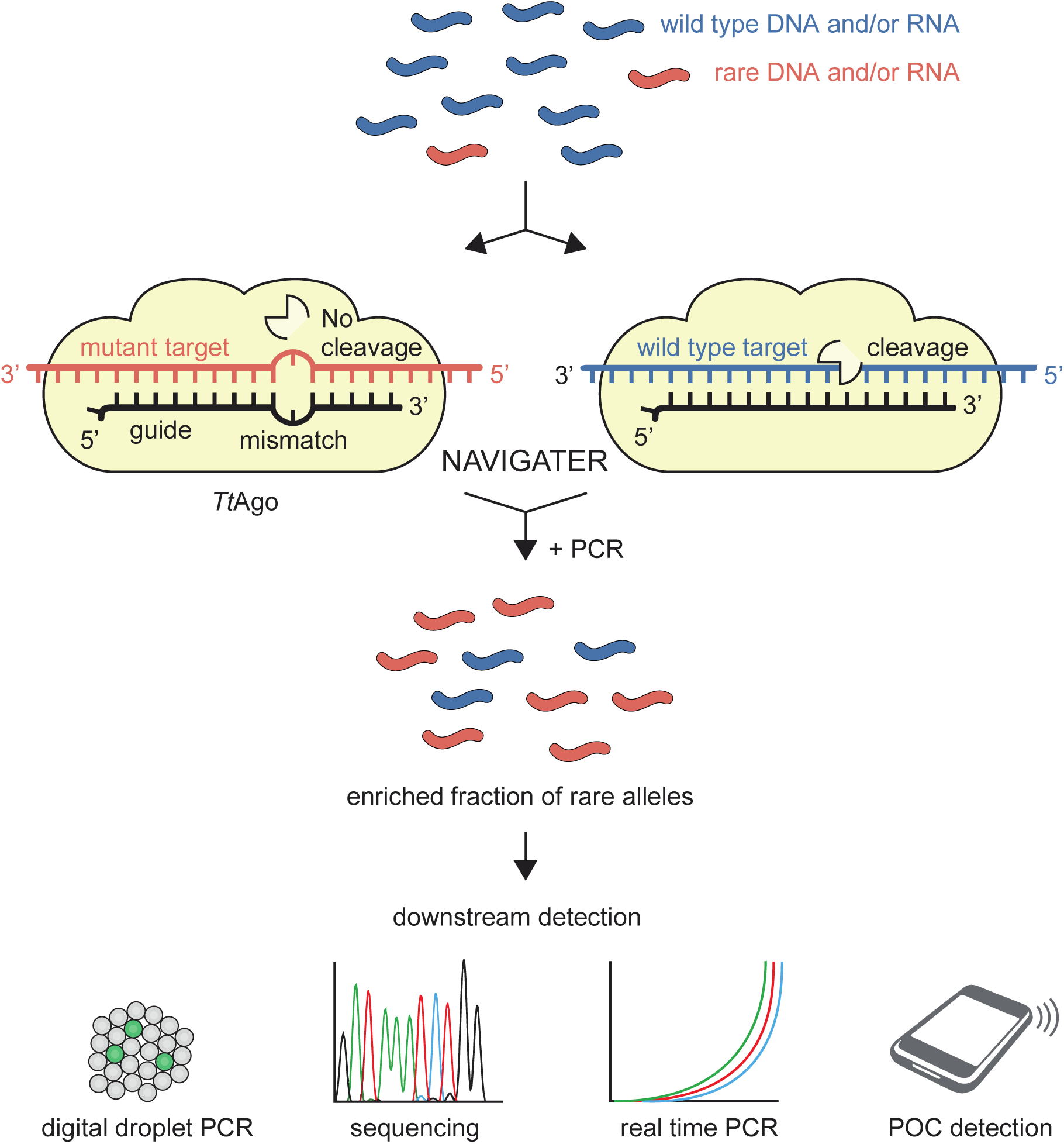
Schematic overview of NAVIGATER. Programmed with a DNA guide, *Tt*Ago specifically cleaves abundant, fully complementary WT alleles (both DNA and RNA, blue) while sparing rare alleles (red) with a single nucleotide mutation. This results in the enrichment of the fraction of disease-related rare nucleic acids, enhancing the sensitivity of downstream detection methods.

## Results

### Betaine, Mg^2+^, and dNTPs enhance *Tt*Ago’s endonucleolytic activity

We first examined *Tt*Ago activity in various buffers (Supplemental Section 1) and selected a buffer that provided the best performance (Buffer 3, Supplemental Table 1). Significantly, we find that betaine, dNTPs, and Mg^2+^ enhance *Tt*Ago’s activity without adversely impacting its single nucleotide specificity. The same buffer is suitable for cleaving both DNA and RNA.

**Table 1:** Comparison of rare allele enrichment methods.

|  | **dCas9-based method<sup>12</sup>** | **Cas9-DASH<sup>10</sup> and Cut-PCR<sup>11</sup>** | **NAVIGATER** | **NaME-Pro<sup>37</sup>** | **COLD-PCR<sup>38</sup>** | **Blocker (PNA or DNA)-PCR<sup>39,40</sup>** |
| --- | --- | --- | --- | --- | --- | --- |
| **Enzyme** | deactivated CRISPR Cas9 (high cost) | CRISPR Cas proteins (low cost) | *TtAgo* (low cost) | DSN (high cost) | N/A* | N/A |
| **Guide/blocker** | ~120 nt sgRNA (high cost) | ~120 nt sgRNA (high cost) | 15/16 nt DNA (low cost) | 20-25 nt DNA (low cost) | N/A | PNA (high cost)<br>DNA (low cost) |
| **PAM site requirement** | Yes | Yes | No | No | No | No |
| **Multi-turnover enzyme** | No | No | Yes | Yes | N/A | N/A |
| **Target** | DNA | DNA | DNA & RNA | DNA | DNA | DNA |
| **Incubation time** | ~7 h | 2 h | <1 h | 22 min | ~2 h | ~2 h |
| **Tight temperature control** | No (37°C) | No (37°C) | No (66-86°C) | Yes** (98°C, 67°C) | Yes (thermal cycling) | Yes (thermal cycling) |
| **Fraction detection limit** | 0.1% with allele-specific qPCR | 0.01% with targeted deep sequencing | 0.01% with XNA-PCR | 0.01% with HRM/Sanger sequencing | 0.1%-0.5% with MALDI-TOF | 0.1%-1% |
| **Enrichment level** | Medium | High | High | High | High | High |
| **Specificity** | Medium | Medium-high | High | Medium | Medium | Medium |
| **Multiplexing capability** | Yes | Yes | Yes | Yes | Yes | Yes |

\* N/A – not applicable
\*\* After incubating at 98°C for 2 minutes, multiple steps, such as maintaining the temperature at 67°C and then opening tube to add DSN are required.

### Single guide-target-mismatches curtail target cleavage

Ago proteins’ cleavage efficiency depends sensitively on the positions of single and dinucleotide mismatches between a guide oligo and target strand^22,23^. Since we require efficient cleavage of wild type (WT) alleles while sparing mutant alleles with a single nucleotide mismatch with gDNA, we examined the effects of guide’s nucleotide mismatch position (MP, counted from the gDNA’s 5’ end) on cleavage efficiency of WT *KRAS, BRAF, EGFR*, and their mutants – both DNA and RNA (Fig. 2 and Supplementary Section 2). Fig. 2a depicts the various gDNAs tested with *KRAS* G12D. *Tt*Ago-mediated cleavage of *KRAS* G12D strand is nearly completely curtailed when gDNA has a nucleotide mismatch at MP7 or MP11-13 (Fig. 2b). Guides designed for enriching *KRAS* G12D also enrich other *KRAS* G12 mutations.

**Fig. 2.**
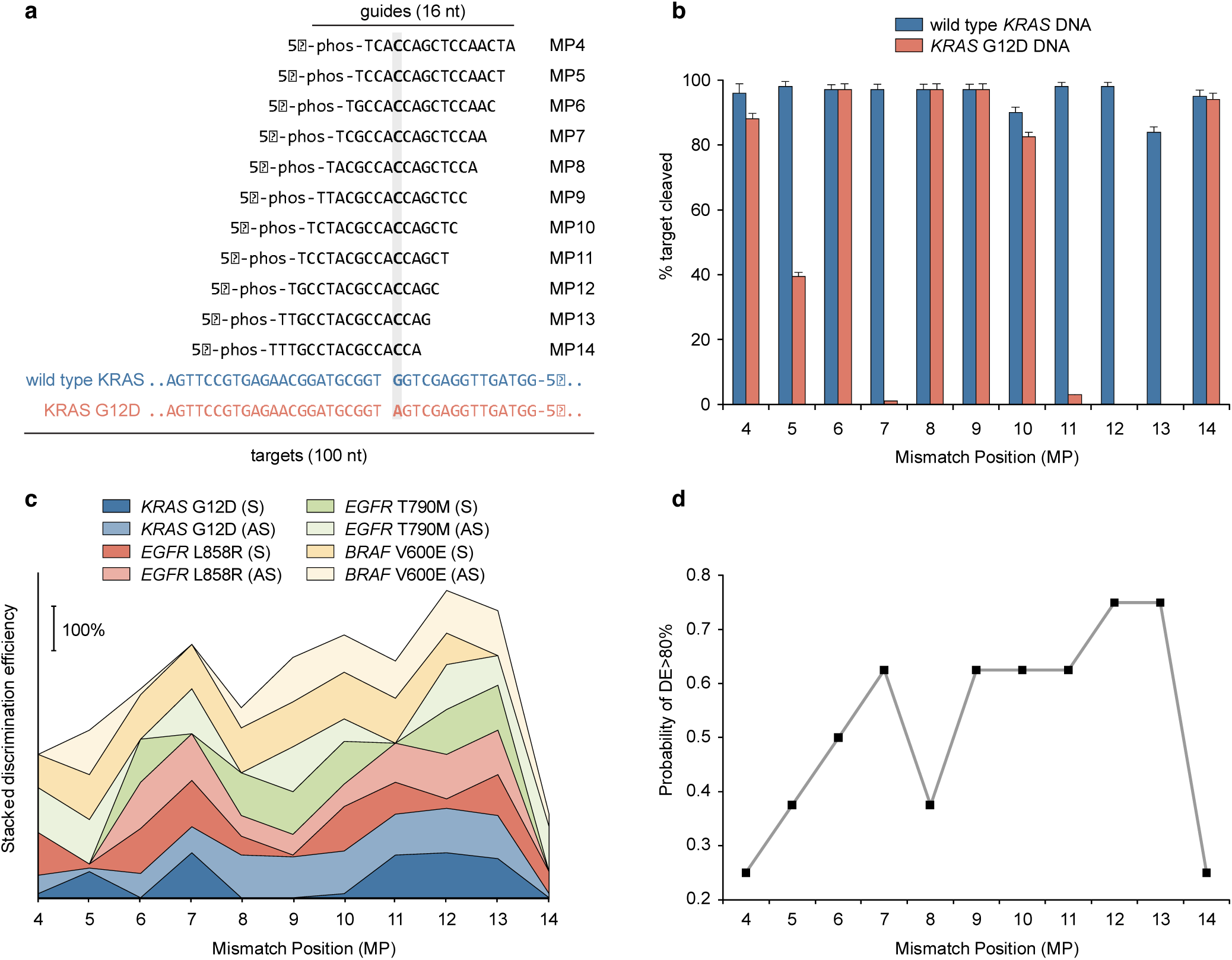
NAVIGATER discrimination efficiency (DE) depends sensitively on the position of the mismatch (MP) between gDNA and allele. **a**, *KRAS* – sense (S) guide and S target sequences. **b**, Cleavage efficiencies of WT *KRAS* DNA (S) and *KRAS* G12D DNA (S) as functions of MP. Error bars show 1 standard deviation, N=3. **c**, Stacked DE for *KRAS* G12D, *EGFR* L858R, *EGFR* T790M, and *BRAF* V600E DNA as a function of MP. AS = Antisense. **d**, Probability of DE >80% as a function of MP for the cases in (c). Experiments were carried out with short guides (15/16 nt) in Buffer 3 at 80 °C. *Tt*Ago: guide: target = 5 : 1 : 1 (1.25 μM : 0.25 μM : 0.25 μM). The notation MP-X indicates the position (X) of the mismatch (MP) between guide oligo and allele counted from the 5’ end of the guide.

To characterize gDNA performance, we define the discrimination efficiency (DE) as the difference in cleavage efficiency of the WT and mutant strand. Fig. 2c depicts DE as a function of MP for *KRAS, EGFR* and *BRAF* DNA and their mutants. Mismatches MP7 and MP9-MP13, located around the cleavage site (g10/g11), yielded the best discrimination (Fig. 2c, d). The optimal MP depends, however, on the allele’s specific sequence. Cleavage of RNA was less tolerant to mismatches than cleavage of DNA. Single mismatches anywhere between MP4-MP11 near completely prevented RNA cleavage (Supplementary Figs. 6e and 8).

Despite being loaded with guides that were fully complementarity to the target, WT sequences were cleaved by *Tt*Ago at variable efficiencies (Fig. 2b and Supplementary Section 2). This suggests that besides the different buffer components, the activity of *Tt*Ago additionally depends on the sequence of the guide and target, which might affect the conformation of the ternary *Tt*Ago-gDNA-DNA and *Tt*Ago-gDNA**-**RNA complexes^24^.

### *Tt*Ago cleaves most specifically with short guides (15/16 nt)

Heterologously-expressed *Tt*Ago is typically purified with gDNAs ranging in length from 13 to 25 nt^17^, whereas *in vitro, Tt*Ago operates with gDNAs ranging in length from 7 to 36 nt^15^. Since little is known on the effect of guide’s length on *Tt*Ago*’s* discrimination efficiency (DE), we examine this effect in our *in-vitro* assay. *Tt*Ago efficiently cleaves WT *KRAS* (sense, S) with complementary guides, ranging in length from 16 to 21 nt at both 70°C and 75°C (Fig. 3a and Supplementary Figs. 11 and 12). Guides of 17-21 nt length with a single nucleotide mismatch at MP12 cleave DNA mutant strands at 75°C, but not at 70°C (Fig. 3a). Cleavage of the mutant allele at 75°C is, however, completely suppressed with a 16 nt guide (Fig. 3a, bottom). We observe similar behavior when a 15 nt guide with a single mismatch at MP13 targets the antisense (AS) *KRAS* strand (Supplementary Fig. 11a-ii). We hypothesize that shorter guides form a less stable complex with off-targets than longer guides, preventing undesired cleavage.

**Fig. 3.**
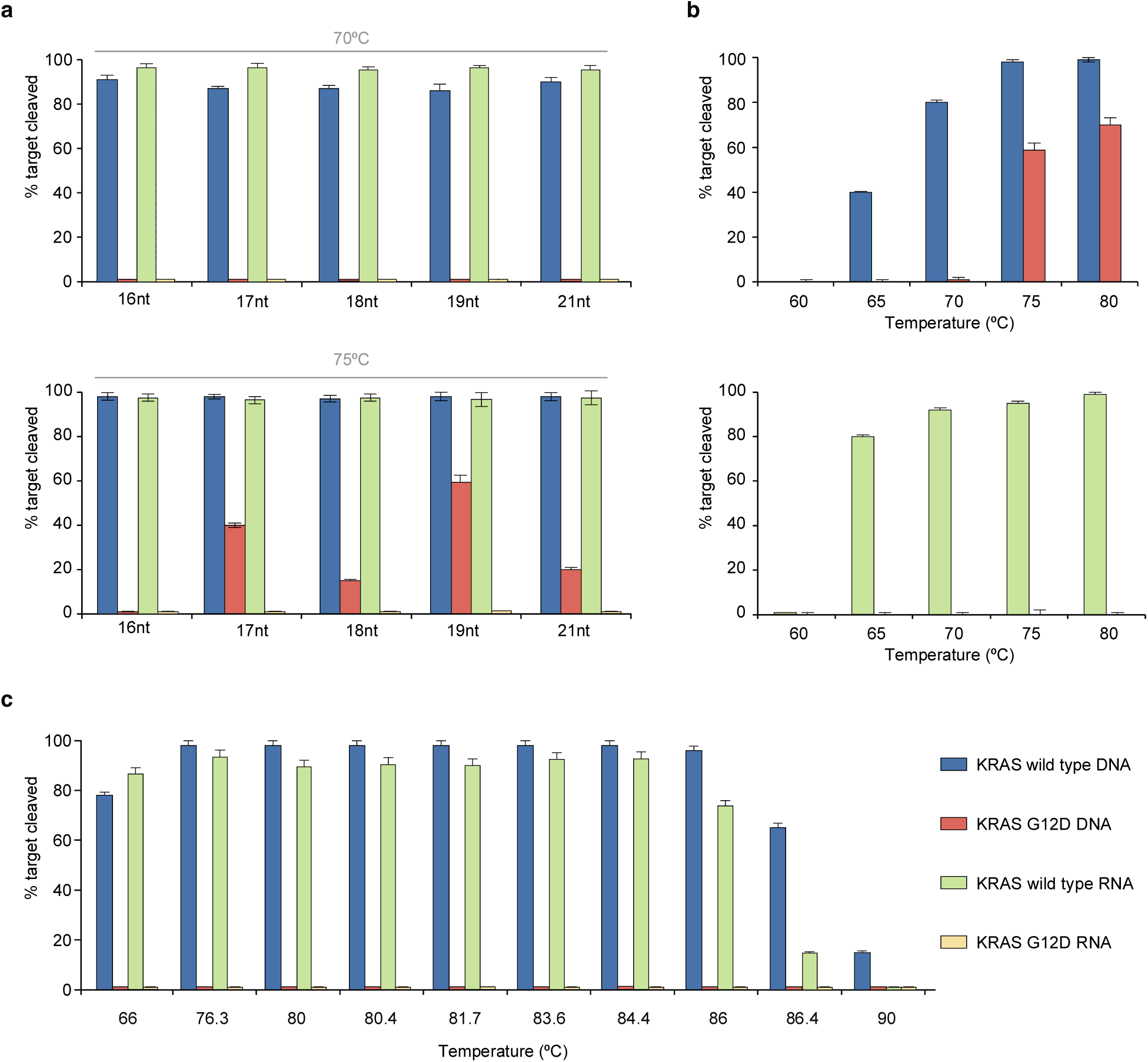
*Tt*Ago cleaves more specifically with short guides. **a**, Cleavage efficiencies as functions of guide length at 70°C (top) and 75°C (bottom). *Tt*Ago in complex with the MP12 guide was used to target either *KRAS* (S) WT or G12D. **b**, Cleavage efficiencies of (top) DNA (S) and (bottom) RNA (S) as functions of temperature with 19 nt long guide (*KRAS*-S (19nt)-MP12). **c**, Cleavage efficiencies of DNA (S) and RNA (S) as functions of temperature with 16 nt long guide (*KRAS*-S (16nt)-MP12). Experiments were carried out in Buffer 3 with *Tt*Ago: guide: target ratio 5: 1: 1. N=3.

In contrast to mutant DNA, an increase in assay temperature does not increase undesired cleavage of mutant RNA (Fig. 3a, b), likely due to differences in the effects of ssDNA and ssRNA on enzyme conformation. In summary, when operating with short guides (15/16 nt), *Tt*Ago efficiently cleaves both WT RNA and DNA targets in the same buffer while avoiding cleavage of mutant alleles at temperatures ranging from 66 to 86°C (Fig. 3c).

### *Tt*Ago efficiently cleaves targeted dsDNA at temperatures above the dsDNA’s melting temperature

Guide-free (apo-) *Tt*Ago can degrade dsDNA via a ‘chopping’ mechanism, autonomously generating functional gDNA^25^. This is, however, a slow process that takes place only when the target DNA is rich in AT (<17% GC)^17^, suggesting that *Tt*Ago lacks helicase activity and depends on dsDNA thermal breathing to enable chopping^13,17,26^. In our assays *Tt*Ago is saturated with gDNAs to suppress this apo activity on dsDNA. *Tt*Ago’s activity at high temperatures provides NAVIGATER with a clear advantage since dsDNA unwinds as the incubation temperature increases.

We determined the optimal temperature at which *Tt*Ago, saturated with guides, efficiently cleaves ds*KRAS* WT while sparing the mutant allele. The estimated melting temperature of 100 bp ds*KRAS* (S strand sequence listed in Fig. 2a) in Buffer 3 is 79.7°C (IDT-OligoAnalyzer)^27^. Consistent with this estimate, very little cleavage of dsDNA takes place at temperatures below 80°C, but *Tt*Ago cleaves dsDNA efficiently at temperatures above 80°C (Fig. 4a-i). Cleavage efficiency increases as the incubation time increases and nearly saturates after about one hour (Figs. 4b). *Tt*Ago efficiently cleaves high GC-content (∼70%) dsDNA at 83°C (Supplementary Fig. 13) due to the high betaine concentration in our buffer^28^.

**Fig. 4.**
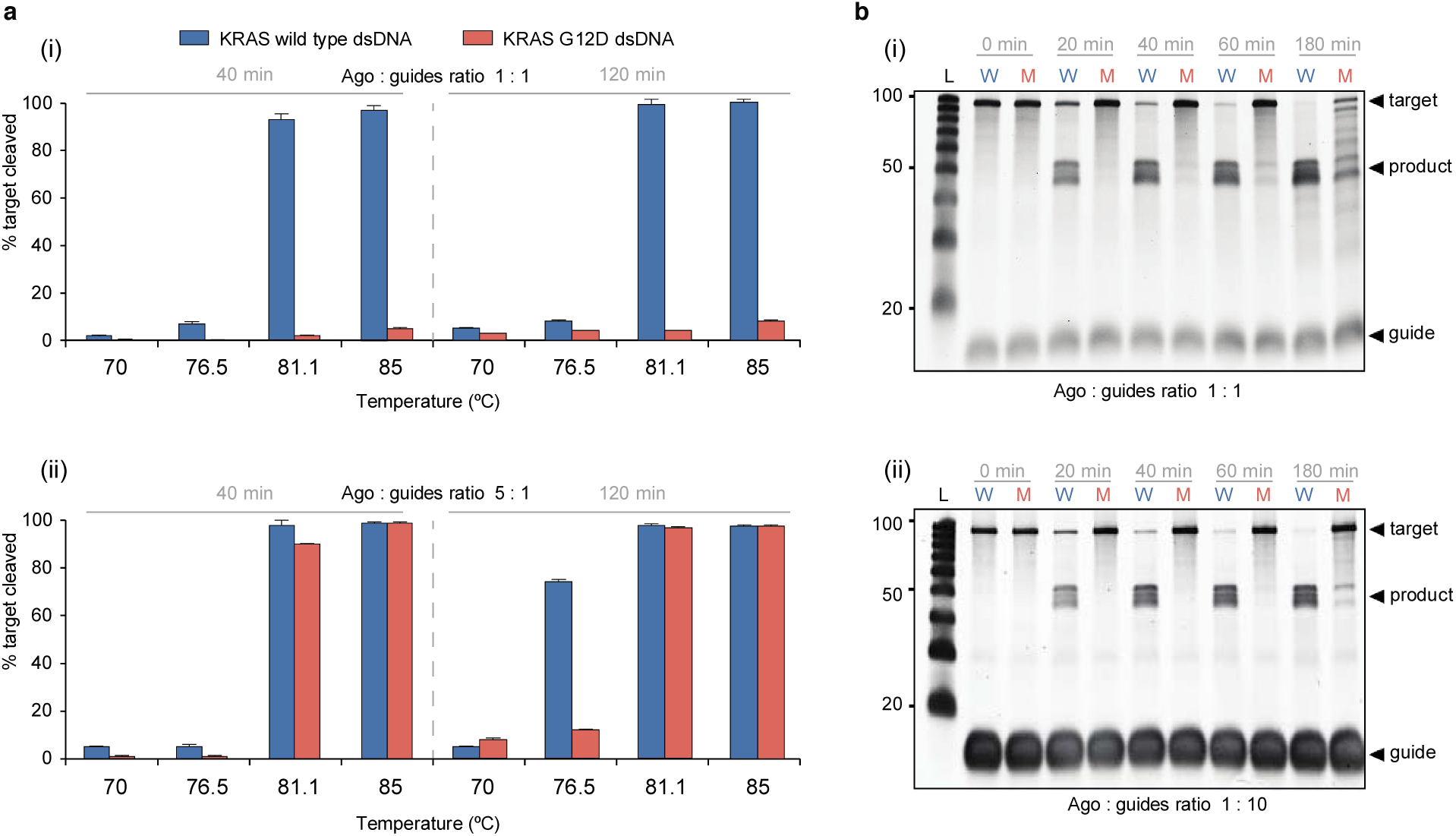
Excess guide concentration provides high dsDNA cleavage discrimination efficiency. **a**, The cleavage efficiencies of *KRAS* WT and *KRAS* G12D dsDNA as functions of temperature: (**i**) *Tt*Ago/guide ratio: 1: 1 (1.25 μM : 1.25 μM), 40 min and 120 min incubation times; (**ii**) *Tt*Ago/guide ratio: 5: 1 (1.25 μM : 0.25 μM), 40 min and 120 min incubation times. **b**, Electropherograms of NAVIGATER’s products as a function of incubation time (83°C): (**i**) *Tt*Ago/guide ratio 1: 1 and (**ii**) *Tt*Ago/guide ratio 1: 10 (1.25 μM : 12.5 μM). All experiments were carried out in Buffer 3 with *KRAS*-S (16nt)-MP12 and *KRAS*-AS (15nt)-MP13 guides. N=3.

### Guide saturation is necessary to avoid off-target cleavage

In the presence of guide concentrations less than the *Tt*Ago concentration, e.g., *Tt*Ago/guides ratio 5:1, non-specific, undesired cleavage of ds mutant alleles takes place (Fig. 4a-ii). This off-target cleaving increases as the incubation time increases (Fig. 4a-ii) and can be attributed to chopping^25^. Since *Tt*Ago binds gDNA with high affinity, excess of gDNAs eliminates undesired chopping (Figs. 4b and Supplementary Fig. 14). Indeed, at a *Tt*Ago/guides ratio of 1:10, *Tt*Ago efficiently cleaves WT *KRAS, BRAF* and *EGFR* dsDNA while sparing the mutant dsDNA harboring point mutations: *KRAS* G12D (Fig. 4b-ii), *KRAS* G12V (Supplementary Fig. 15), *BRAF* V600E, *EGFR* L858R, and *EGFR* T790M (Supplementary Fig. 16), and deletion mutations in *EGFR* exon 19 (Supplementary Fig. 17). For optimal discrimination between ds WT and mutant alleles, it is necessary to saturate *Tt*Ago with guides and keep the incubation time under an hour. Furthermore, tight *Tt*Ago/guide complexes can be formed by pre-incubation on ice for 3 min (Supplementary Fig. 15).

### *Tt*Ago is more specific than CRISPR/Cas9-based dsDNA cleavage

The recently-developed assays Cas9-DASH^10^ (**D**epletion of **A**bundant **S**equences by **H**ybridization) and CUT-PCR^11^ (**<u>C</u>**RISPR-mediated **<u>U</u>**ltra-sensitive detection of **<u>T</u>**arget DNA-PCR) take advantage of the low tolerance of CRISPR/Cas proteins to mismatches at the PAM recognition site to discriminate mutant from WT alleles. We examine the discrimination efficiency of CRISPR/Cas9 with a previously reported guide RNA^10-12^ (Supplementary Fig. 18a). CRISPR/Cas9 nonspecifically cleaves both ds WT and mutant alleles harboring *KRAS* G12D and G12V mutations (Supplementary Fig. 18b, c). Consistent with previous reports^11,29,30^, we suspect that these non-specific cleavages are caused by non-canonical PAM recognition. CRISPR/Cas9 also failed to differentiate between ds WT and mutant alleles harboring *EGFR* L858R mutation (Supplementary Fig. 18b, c), presumably because of the presence of a PAM site in *EGFR* L858R allele and a non-canonical PAM in the WT allele (Supplementary Fig. 18a), which complicates the design of a guide to specifically cleave the WT while sparing the mutant. In contrast, CRISPR/Cas9 specifically cleaved ds WT *EGFR* while sparing the ds mutant allele harboring the deletion mutation E746-A750 del (1). In summary, CRISPR/Cas9 shows a lower discrimination efficiency compared to our *Tt*Ago system.

### NAVIGATER increases the sensitivity of downstream rare allele detection

In recent years, there has been a rapidly increasing interest in applying liquid biopsy to detect cell-free circulating nucleic acids associated with somatic mutations for, among other things, cancer diagnostics, tumor genotyping, and monitoring susceptibility to targeted therapies. Liquid biopsy is attractive since it is minimally invasive and relatively inexpensive. Detection of mutant alleles is, however, challenging due to their very low concentrations in liquid biopsy samples among the background of highly abundant WT alleles that differ from mutant alleles by as little as a single nucleotide. To improve the sensitivity and specificity of detecting rare alleles that contain valuable diagnostic and therapeutic clues, it is necessary to remove and/or suppress the amplification of WT allelesxs^31-33^. NAVIGATER meets this challenge by selectively and controllably degrading WT alleles in the sample, thereby increasing the fraction of mutant alleles. We demonstrate here that NAVIGATER increases sensitivity of downstream mutation detection methods such as gel electrophoresis, ddPCR^18^, PNA-PCR^19^, PNA-LAMP^20^, Sanger sequencing, and XNA-PCR^21^; and enables multiplexed enrichment. To demonstrate NAVIGATER’s potential clinical utility, we enriched blood samples from pancreatic cancer patients (Supplementary Table 2) that were previously analyzed with RainDance ddPCR. These samples were pre-amplified by PCR to increase WT and mutant *KRAS* total content before enrichment.

#### Gel electrophoresis

(Supplementary Fig. 19): We subjected the enrichment assay products to gel eIectrophoresis. In the absence of enrichment (control), the bands at 80 bp (*KRAS*) on the electropherogram are dark. After 40 minutes of *Tt*Ago enrichment, these bands faded, indicating a reduction of *KRAS* WT alleles. After 2 hours enrichment, all the bands at 80 bp, except that of patient P6, have essentially disappeared, suggesting that most WT alleles have been cleaved. The presence of an 80 bp band in the P6 lane is attributed to the relatively high (20%) mutant allele frequency (MAF) that is not susceptible to cleavage. We also PCR amplified the products from a 2-hour NAVIGATER treatment, and subjected the amplicons to a second NAVIGATER (2h). The columns P3, P4, and P6 feature darker bands than P1, P2 and P5, indicating the presence of mutant alleles in samples P3, P4, and P6 (Supplementary Fig. 19) and demonstrating that NAVIGATER renders observable otherwise undetectable mutant alleles.

#### Droplet Digital PCR (ddPCR)

(Supplementary Fig. 20): To quantify our enrichment assay products, we subjected them to ddPCR. The detection limit of ddPCR is controlled by the number of amplifiable nucleic acids in the sample, which must be a small fraction of the number of ddPCR droplets. The large number of WT alleles in the sample limits the number of pre-ddPCR amplification cycles that can be carried out to increase rare alleles’ concentration. Since NAVIGATER drastically reduces the number of WT alleles in the sample, it enables one to increase the number of pre-amplification cycles, increasing the number of mutant allele copies, and thus, the ddPCR sensitivity. When operating with a mixture of WT and mutant allele, NAVIGATER products include: residual uncleaved WT (*N*_*WT*_), mutant (*N*_*M*_), and WT-mutant hybrids (*N*_*H*_) alleles. Hybrid alleles form during re-hybridization of an ss WT with an ss mutant allele. The MAF is *f*_*M*_*=(N*_*M*_+*½N*_*H*_*)/(N*_*WT*_+*N*_*M*_+*N*_*H*_).

We carried out ddPCR on un-enriched (control), once-enriched, and twice-enriched samples (Supplementary Fig. 20b), increasing MAF significantly (Fig. 5a). For example, the MAF increased from 0.5% in the un-enriched P5 (G12D) sample to ∼30% in the twice-enriched sample. This represents an ∼60 fold increase in the fraction of droplets containing the mutant allele (Fig. 5b). The same assay also enriched G12R, increasing MAF from 3% to ∼66% in sample P3 and G12V, increasing MAF from 5% to ∼68% in sample P4 (Fig. 5b).

**Fig. 5.**
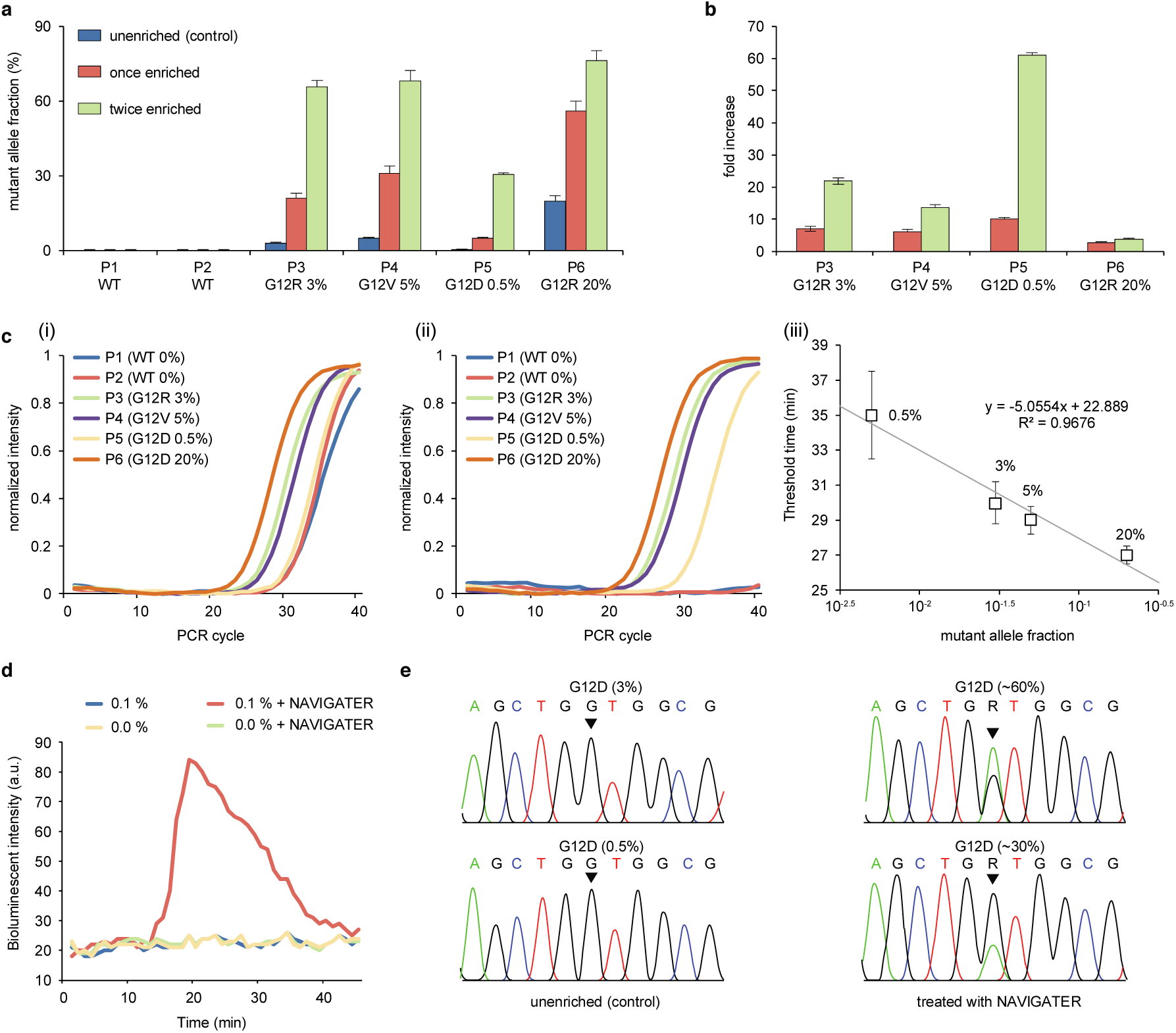
NAVIGATER enhances the sensitivity of downstream detection methods. **a, b**, ddPCR of samples from pancreatic cancer patients containing *KRAS* mutants (Supplementary Table 2): **a**, Fraction of droplets reporting mutant alleles in the absence of enrichment (control), once enriched, and twice enriched. **b**, Increase in mutant allele frequency after NAVIGATER enrichment. **c**, PNA-PCR’s amplification curves of pancreatic cancer patients’ samples before (**i**) and after (**ii**) NAVIGATER, and amplification threshold time as a function of MAF (**iii**). **d**, PNA-LAMP of simulated RNA samples before and after NAVIGATER carried out with a minimally-instrumented, electricity-free Smart-Connected Cup (SCC)^34^. **e**, Sanger sequencing before and after NAVIGATER when detecting simulated RNA samples. All the controls were pre-processed in the absence of *Tt*Ago. N=3

#### PNA-PCR

PNA-PCR engages a sequence-specific PNA blocker that binds to WT alleles, suppressing WT amplification and providing a limit of detection of ∼1% MAF ^19^. To demonstrate NAVIGATER’s utility, we compared the performance of PNA-PCR when processing pancreatic cancer patient samples (Supplementary Table 2) before and after NAVIGATER (Fig. 5c). Before enrichment, PNA-PCR real-time amplification curves in the order of appearance are P6, P4, and P3, as expected. Samples P1 (MAF=0%), P2 (MAF=0%), and P5 (MAF=0.5%) nearly overlap, consistent with a detection limit of ∼1%^19^. Enrichment (Fig. 5c-ii) significantly increases the threshold times of samples P1 and P2, revealing the presence of mutant alleles in sample P5. PNA-PCR combined with NAVIGATER provides the linear relationship *T*_*½*_*= 22.9-5 log(MAF)* between threshold time (the time it takes the amplification curve to reach half its saturation value) and MAF (Fig. 5c-iii), allowing one to estimate mutant allele concentration. The data suggests that NAVIGATER can improve PCR-PNA limit of detection to below 0.1%.

#### PNA-LAMP

Genotyping with PNA blocking oligos can be combined with the isothermal amplification LAMP^20^. To demonstrate the feasibility of genotyping at the point of care and in resource-poor settings, we use a minimally-instrumented, electricity-free Smart-Connected Cup (SCC)^34^ with smartphone and bioluminescent dye-based detection to incubate PNA-LAMP and detect reaction products. To demonstrate that we can also detect RNA alleles, we used simulated samples comprised of mixtures of WT *KRAS* mRNA and *KRAS-*G12D mRNA. In the absence of pre-enrichment, SSC is unable to detect the presence of 0.1% *KRAS-*G12D mRNA whereas with pre-enrichment 0.1% *KRAS-*G12D mRNA is readily detectable (Fig. 5d).

#### Sanger Sequencing

In the absence of enrichment, Sanger sequencers detect >5% MAF^35^. The Sanger sequencer failed to detect the presence of MAF *∼*3% and 0.5% *KRAS-G12D* mRNA in our un-enriched samples, but readily detected these mutant alleles following NAVIGATER enrichment (Fig. 5e).

#### Multiplexed enrichments

We carried out triplex NAVIGATER and compared it with triplex Cas9 - based DASH (Cas9-DASH)^10^. Our experiment demonstrates that NAVIGATER can operate as a multiplexed assay, enriching multiple mutant alleles in a single sample; it is more specific than CRISPR/Cas9’s PAM site recognition-based enrichment; and it can be combined with clamped assay XNA-PCR^21^ that is similar to PNA-PCR to significantly improve XNA-PCR sensitivity (Supplementary Section 7).

### NAVIGATER enables detection of low-frequency (<0.2%) mutant alleles in blood samples of pancreatic cancer patients

Blood samples were collected from 18 pancreatic cancer patients and 4 healthy control donors, processed to plasma, and banked at −80°C. All patients had clinical NGS performed on a routinely collected tissue sample, 14 tested positive and 4 tested negative for *KRAS* G12D, V, or R mutations (Supplementary Table 4). Below, we compare our blood test results to the tissue data (gold standard). Pre-amplification followed by *KRAS* G12D/V/R mutation detection by RainDance ddPCR was performed on cfDNA extracted from 0.75 mL of patient plasma. An equivalent (blinded) cfDNA sample was pre-amplified for the *KRAS* G12 region (DiaCarta, Inc^21^). Equal volumes of the resultant amplicons were subjected in duplicate to (1) XNA-PCR without pre-enrichment, (2) cas9-DASH followed by XNA-PCR, and (3) single NAVIGATER (once-NAVIGATER) followed by XNA-PCR. These results were compared to tissue NGS prior to subsequent testing with two NAVIGATER enrichment steps (twice-NAVIGATER). The XNA-PCR test was deemed positive when the threshold number of cycles (Ct) was less than a cutoff (Ct_C_) - the average Ct minus 3SD (95% confidence level) of XNA-PCR of standard (Horizon) WT *KRAS* allele samples (in the absence of mutant alleles) at DNA concentrations greater than in our clinical samples (Supplementary Section 8).

XNA-PCR without pre-enrichment identified 1/14 samples as positive and 1/14 as inconclusive (only one of the duplicates tested positive). Cas9-DASH XNA-PCR identified 3/14 as positive; the Ct values of other samples were clustered together (Supplementary Fig. 23b) complicating discrimination between positives and negatives. Once-NAVIGATER followed by XNA-PCR identified 6/14 samples as positive; twice-NAVIGATER followed by XNA-PCR identified 9/14 as positive (Fig. 6b, c, e). ddPCR identified 8/14 as positive (Fig. 6b, c). The G12D/V/R tissue negative samples and healthy controls are all negative in these tests. The sensitivity, specificity, positive predictive value, negative predictive value, and concordance with tissue NGS genotyping for *KRAS* G12 mutations were, respectively, 64%, 100%, 100%, 62%, 77% for twice-NAVIGATER followed by XNA-PCR, and 57%, 100%, 100%, 57%, 73% for ddPCR (Fig. 6b). The receiver operating characteristic (ROC, Fig. 6d) curve for twice-NAVIGATER followed by XNA-PCR has an area under the ROC curve (AUC) of 0.88 (95% CI: 0.67 - 0.98), indicating that with an optimal Ct_C_, our assay has 79% sensitivity and 100% specificity.

NAVIGATER XNA-PCR identified 3 positives that were undetectable by ddPCR, while ddPCR identified 2 positives that were undetectable by NAVIGATER XNA-PCR (Fig 6e). All positive twice-NAVIGATER-enriched samples were subjected to Sanger sequencing to verify that the *KRAS* G12 variants were identical to those detected in tissue biopsy (Supplementary Fig. 24). Since a few samples had single copies of mutant allele (based on ddPCR, Supplementary Table 4), we attribute the discordance between NAVIGATER XNA-PCR and ddPCR to sampling bias, resulting in some of the samples having no targets. In summary, NAVIGATER XNA-PCR (AUC= 0.85, 95% CI: 0.64-0.96 for once-NAVIGATER, and 0.88 for twice-NAVIGATER) is effective at detecting the presence of rare mutant alleles and outperforms XNA-PCR without pre-enrichment (AUC=0.65, 95% CI: 0.42-0.84) and Cas9-DASH XNA-PCR (AUC=0.83, 95% CI: 0.62-0.96) (Supplementary Fig. 23), consistent with our earlier observations that Cas9 is less specific than *Tt*Ago in sparing mutant alleles.

## Discussion

Liquid biopsy is a simple, minimally invasive, rapidly developing diagnostic method to analyze cell-free nucleic acid fragments in body fluids and obtain critical information on patient health and disease status. Currently, Liquid biopsy helps personalize and monitor treatment for patients with advanced cancer, but the sensitivity of available tests is insufficient for patients with early stage disease^33^ or for cancer screening. Detection of alleles that contain critical clinical information is challenging since they are present at very low concentrations among abundant background of nucleic acids that differ from alleles of interest by as little as a single nucleotide.

Here, we report on a novel enrichment method (NAVIGATER) for rare alleles that uses *Tt*Ago. *Tt*Ago is programmed with short ssDNA guides to specifically cleave guide-complementary alleles, *both* DNA and RNA, and stringently discriminate against off-targets with single nucleotide precision. Sequence mismatches between guide and off-targets reduce hybridization affinity and cleavage activity by sterically hindering formation of a cleavage-compatible state^15,16^. *Tt*Ago’s activity and discrimination efficiency depend sensitively on the (i) position of the mismatched pair along the guide, (ii) buffer composition, (iii) guide length, (iv) Ago/guides ratio, (v) incubation temperature and time, and (vi) target sequence. *Tt*Ago appears to discriminate best between target and off-target in the presence of a mismatch at or around the cleavage site located between guide nucleotides 10 and 11. Optimally, the buffer should contain ≥ 8 mM [Mg^2+^], ≥ 0.8 M betaine, and 1.4 mM dNTPs. The ssDNA guides should be 15-16nt in length with their concentration exceeding *Tt*Ago’s concentration; and the incubation temperature should exceed the target dsDNA melting temperature. NAVIGATER is amenable to multiplexing and can concurrently enrich multiple mutant alleles in a single sample while operating with guides with different sequences.

NAVIGATER successfully enriches the fraction of cancer biomarkers such as *KRAS, BRAF*, and *EGFR* mutants in various samples. For example, NAVIGATER increased *KRAS* G12D fraction from 0.5% to 30% (60 fold) in a blood sample from a pancreatic cancer patient. The presence of 0.5% *KRAS* G12D could not be detected with Sanger sequencing or PNA-PCR. However after NAVIGATER pre-processing, both the Sanger sequencer and PNA-PCR readily identified the presence of *KRAS* G12D. Additionally, NAVIGATER combined with PNA-LAMP detects low fraction (0.1%) mutant RNA alleles, enabling genotyping at the point of care and in resource-poor settings. NAVIGATER improves the detection limit of XNA-PCR by more than 10 fold, enabling detection of rare alleles with frequencies as low as 0.01%. NAVIGATER combined with XNA-PCR exhibits higher sensitivity for detecting KRAS mutants in Pancreatic cancer patients than XNA-PCR alone and Cas9-DASH XNA-PCR.

NAVIGATER differs from previously reported rare allele enrichment methods%^10-12,36-40^ in several important ways (Table 1). First, NAVIGATER is versatile. In contrast to CRISPR-Cas9^10-12^ and restriction enzymes^36^, *Tt*Ago does not require PAM motif or a specific recognition site. A gDNA can be designed to direct *Tt*Ago to cleave any desired target. Second, *Tt*Ago is a multi-turnover enzyme^17^: a single *Tt*Ago-guide complex can cleave multiple targets. In contrast, CRISPR-Cas9 is a single turnover nuclease^41^. Third, whereas CRISPR-Cas9 exclusively cleaves DNA, *Tt*Ago cleaves both DNA and RNA targets with single nucleotide precision. In fact, NAVIGATER can enrich for both rare DNA alleles and their associated exosomal RNAs^42^ in the same assay, further increasing sensitivity. Fourth, *Tt*Ago is robust as it operates over a broad temperature range (66 - 86°C) and unlike PCR-based enrichment methods, such as COLD-PCR^38^ and blocker-PCR^39,40^, does not require tight temperature control. Moreover, as we have shown, NAVIGATER can complement PCR-based enrichment methods. Fifth, *Tt*Ago uses DNA guides rather than RNA guides, which reduce cost and increase assay stability. Sixth, *Tt*Ago is more specific than thermostable duplex-specific nuclease (DSN)^37^. Since DSN non-specifically cleaves all dsDNA, DSN-based assays require tight controls of probe concentration and temperature to avoid non-specific hybridization and cleavage of the rare nucleic acids of interest. Most importantly, as we have demonstrated, NAVIGATER is compatible with many downstream genotyping analysis methods such as ddPCR, PNA-PCR, XNA-PCR, and sequencing. Last but not least, NAVIGATER can operate with isothermal amplification methods such as LAMP, enabling integration of enrichment with genotyping for use in resource poor settings.

Our study indicates that *Tt*Ago-based enrichment has the potential to significantly increase the clinical utility of liquid biopsy and enable, in combination with downstream detection methods (including NGS), detect low concentrations of mutant alleles indicative of various types of cancer. NAVIGATER could be leveraged to help detect other rare allele related to such as genetic disorders in fetal DNA and drug resistant bacteria, and help recognize the minor populations of interest in biological studies.

## Acknowledgements

Dr. Robert M. Greenberg helped with PAGE electrophoresis. Dr. Jennifer E. Phillips-Cremins provided us with access to gel imager. Dr. Changchun Liu provided helpful comments early in this project. Stephanie Yee and Taylor Black assisted with ddPCR. Yaguang Fan assisted with statistical analysis. This work is supported by the NIH grant NCI 1R21CA227056-01 to the University of Pennsylvania, and by grants from the Netherlands Organization of Scientific Research (NWO-ECHO 711013002 and NWO-TOP 714015001) to J.v.d.O.

## Author contributions

J.S. and H.H.B. conceived the project and designed the experiments. J.W.H. expressed and purified *Tt*Ago protein. J.S., J.W.H., J.P. and L.T.A. carried out the experiments. M.G.M. and J.C. assisted, respectively, with PNA-PCR and XNA-PCR. N.B. and J.E.T. extracted cfDNA from patients’ blood and quantified *KRAS* G12 mutations with ddPCR. J.E.T. extracted RNA from cell lines. N.B. and J.E.T. assisted with ddPCR experiments. M.S. and J.M. cultured cell lines. E.C. supervised N.B., J.E.T., M.S., and J.M. and advised on patient samples and ddPCR experiments. J.S., J.W.H., M.G.M., J.v.d.O., and H.H.B. analyzed the data and wrote the manuscript. All authors read and commented on the manuscript.

## Competing interests

University of Pennsylvania and Wageningen University have applied for a patent on NAVIGATER with J.S., J.W.H., M.G.M., J.v.d.O., and H.H.B. listed as co-inventors.

## Human Subjects

This study was approved by Penn Institutional Review Board (IRB PROTOCOL #: 822028)

