## Supplementary Information for "Highly Specific Enrichment of Rare Nucleic Acids using *Thermus Thermophilus* Argonaute with Applications in Cancer Diagnostics" (from another version of this preprint: version 3, 2019-01-15)

### Supplementary Section 1: The effect of various buffer compositions on *Tt*Ago cleavage

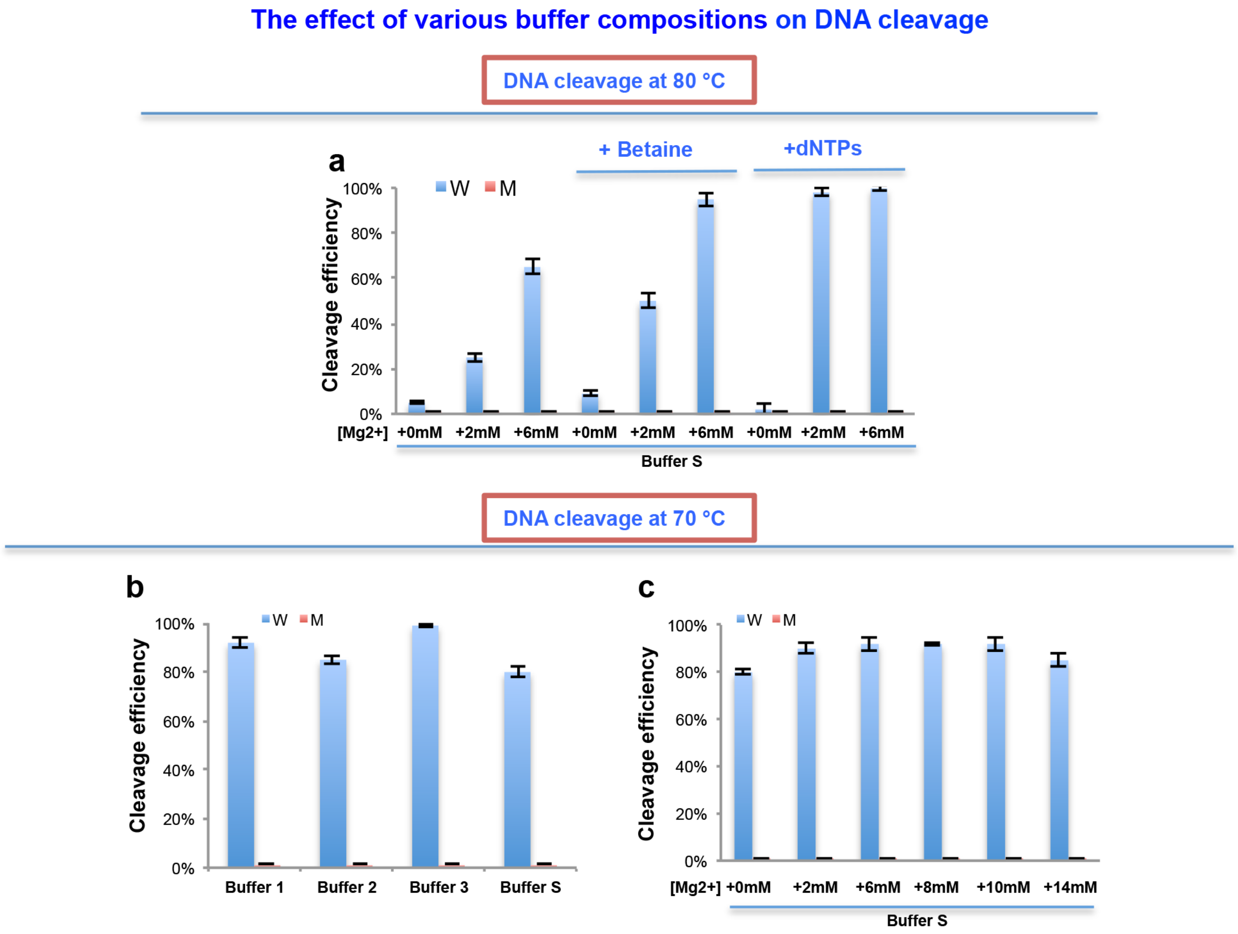


**Supplementary Figure 1: Betaine, $Mg^{2+}$, and dNTPs enhance *Tt*Ago DNA cleavage.** DNA cleavage efficiency as a function of (**a**) added [$Mg^{2+}$] in Buffer S in the absence and presence of betaine (0.8 M) or dNTPs (1.4 mM) at 80 °C; (**b**) Buffer composition (**Table 1**) at 70 °C; and (**c**) added [$Mg^{2+}$] in Buffer S at 70 °C. All experiments were carried out with *KRAS* Sense (S) strand and 16 nt *KRAS*-S MP12 guide. Incubation time 20 min. *Tt*Ago: guide: target =1: 0.2: 0.2. N=3.

### The effect of various buffer compositions on DNA cleavage

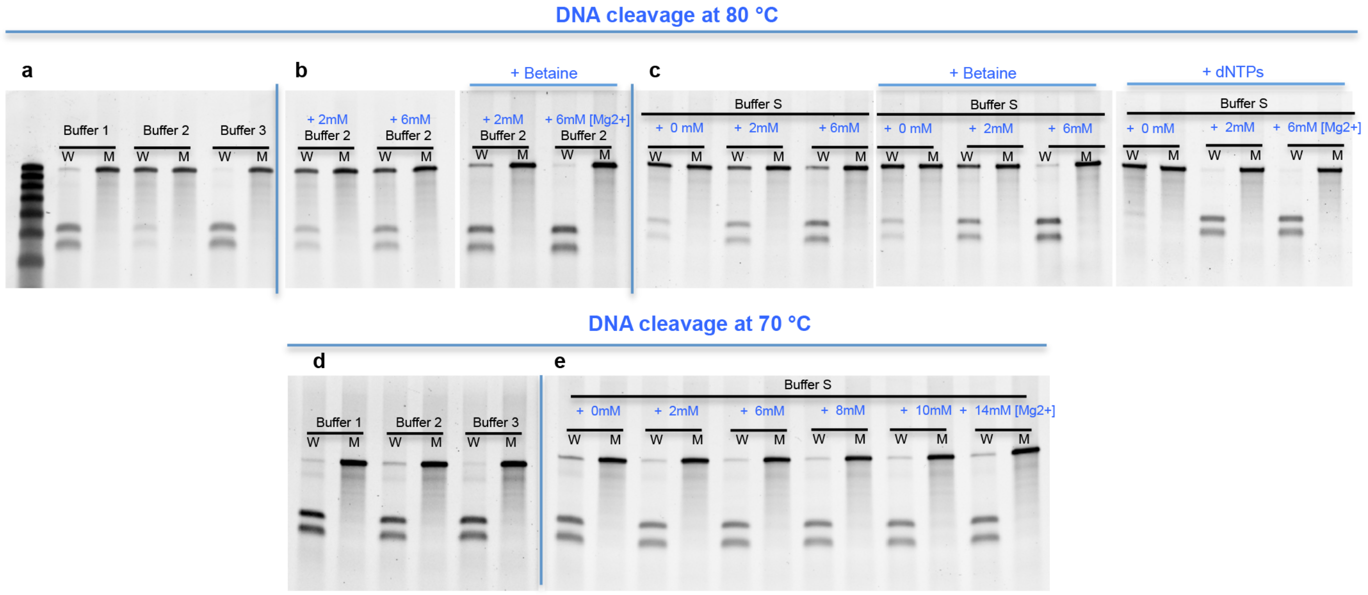


**Supplementary Figure 2. Electropherograms of *Tt*Ago cleavage products of WT *KRAS* (W) and *KRAS* G12D (M) sense ssDNA after 20 min incubation in Buffers 1, 2, 3, and S (Table 1) in the presence and absence of additives.** (**a**) Buffers 1, 2, 3, and S. 80°C, no additives. (**b**) Buffer 2. Various [$Mg^{2+}$] in the absence and presence of betaine (0.8 M), 80°C. (**c**) Buffer S. Various [$Mg^{2+}$] in the absence and presence of betaine (0.8 M) or dNTPs (1.4 mM), 80°C. (**d**) Buffers 1, 2, 3. 70°C, no additives. (**e**) Buffer S. Various [$Mg^{2+}$], 70°C. *Tt*Ago: guide: target =1: 0.2: 0.2. N=3.

**The effect of various buffer compositions on RNA cleavage**

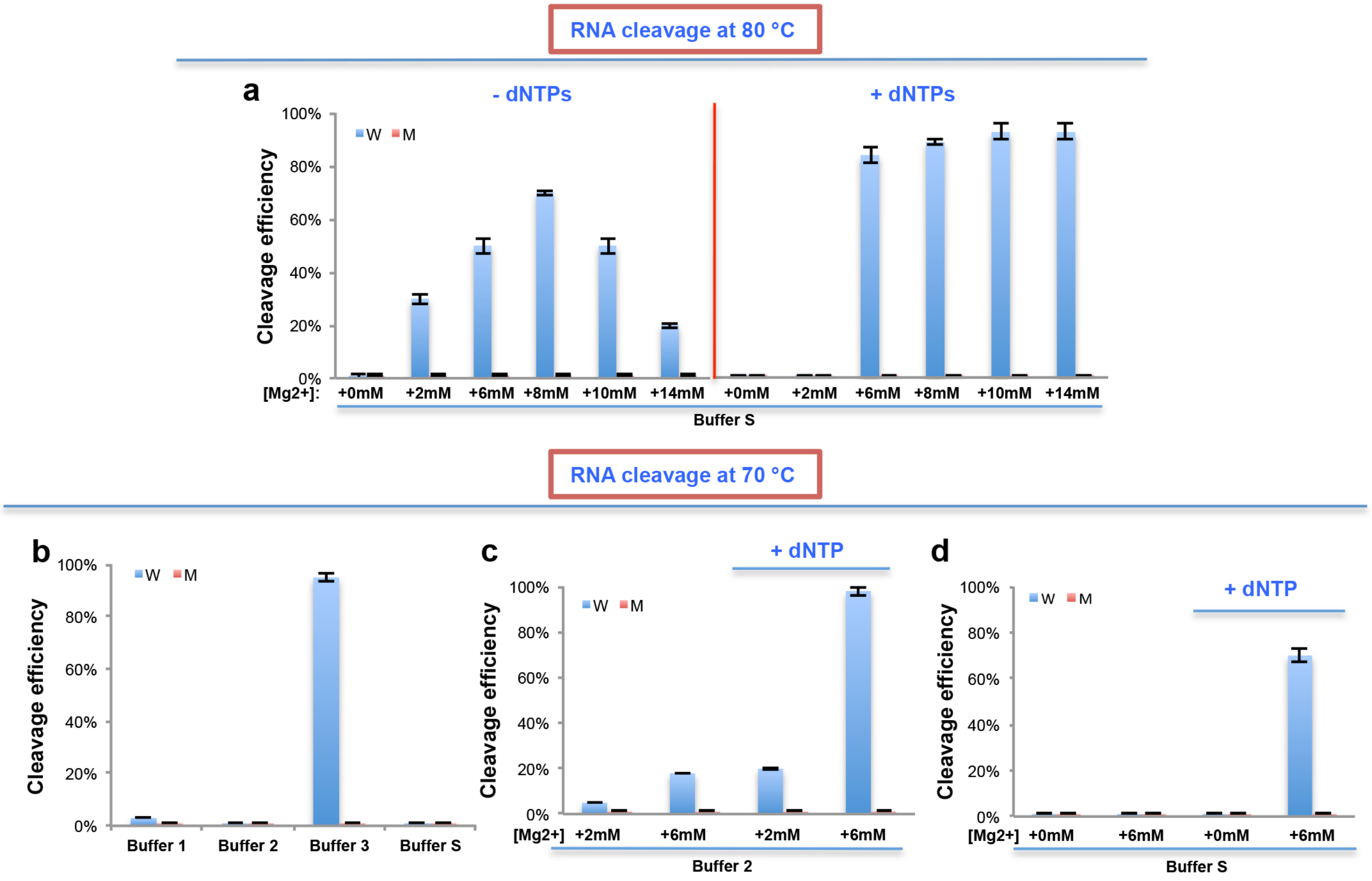


**Supplementary Figure 3: Betaine, $Mg^{2+}$, and dNTPs enhance *Tt*Ago RNA cleavage.** RNA cleavage efficiency as a function of (**a**) added [$Mg^{2+}$] in Buffer S in the absence and presence of dNTPs (1.4 mM) ) at 80 $^{o}$C; (**b**) buffer composition (**Table 1**) at 70 °C; (**c**) added [$Mg^{2+}$] in Buffer 2 in the absence and presence of dNTPs (1.4 mM) at 70°C; and (**d**) added [$Mg^{2+}$] in Buffer S in the absence and presence of dNTPs (1.4 mM) at 70 °C. All experiments were carried out with *KRAS* Sense (S) strand and 16 nt *KRAS*-S MP12 guide. Incubation time 20 min. *Tt*Ago: guide: target =1: 0.2: 0.2. N=3.

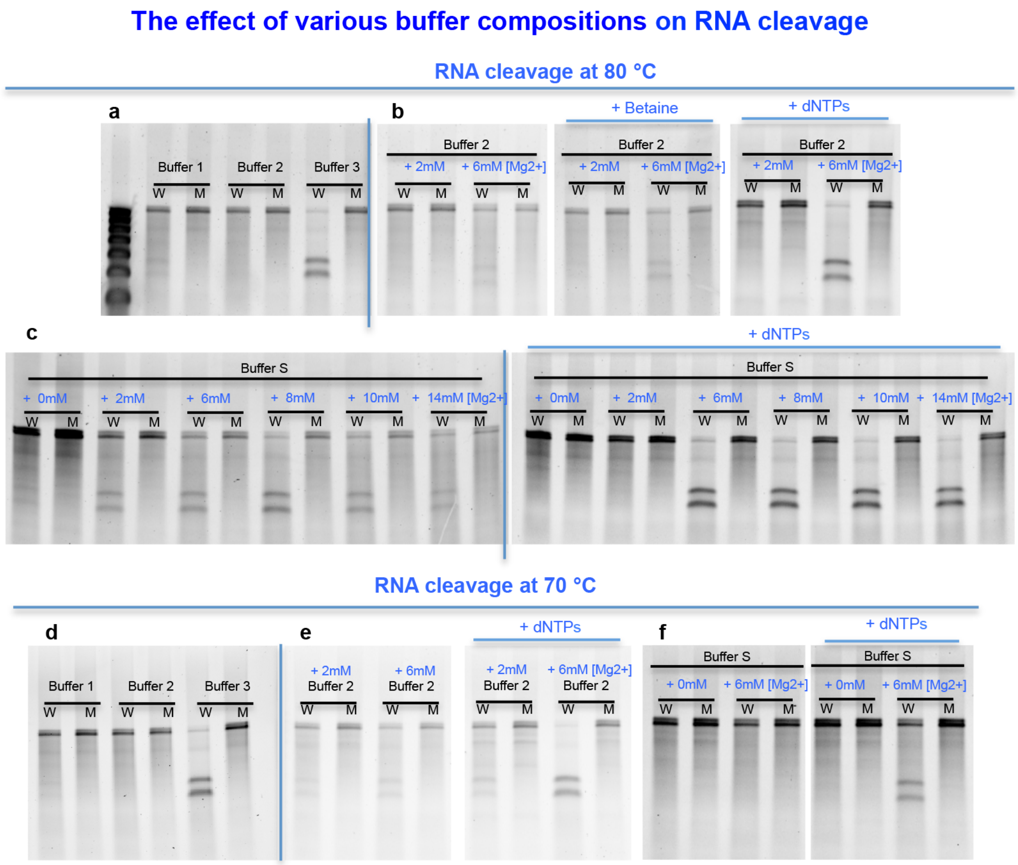


**Supplementary Figure 4. Electropherograms of *Tt*Ago cleavage products of WT *KRAS* (W) and *KRAS* G12D (M) RNA after 20 min incubation in Buffers 1, 2, 3, and S (Table 1) in the presence and absence of additives.** (**a**) Buffers 1, 2, and 3. 80°C, No additives. (**b**) Buffer 2. Various [$Mg^{2+}$] in the absence and presence of betaine (0.8 M) or dNTPs (1.4mM), 80°C. (**c**) Buffer S. Various [$Mg^{2+}$] in the absence and presence of dNTPs (1.4 mM), 80°C. (**d**) Buffers 1, 2, and 3. 70°C, No additives. (**e**) Buffer 2. Various [$Mg^{2+}$] in the absence and presence of dNTPs (1.4 mM), 70°C. (**f**) Buffers S. Various [$Mg^{2+}$] in the absence and presence of dNTPs (1.4 mM), 70 °C. *Tt*Ago: guide: target =1: 0.2: 0.2. N=3.

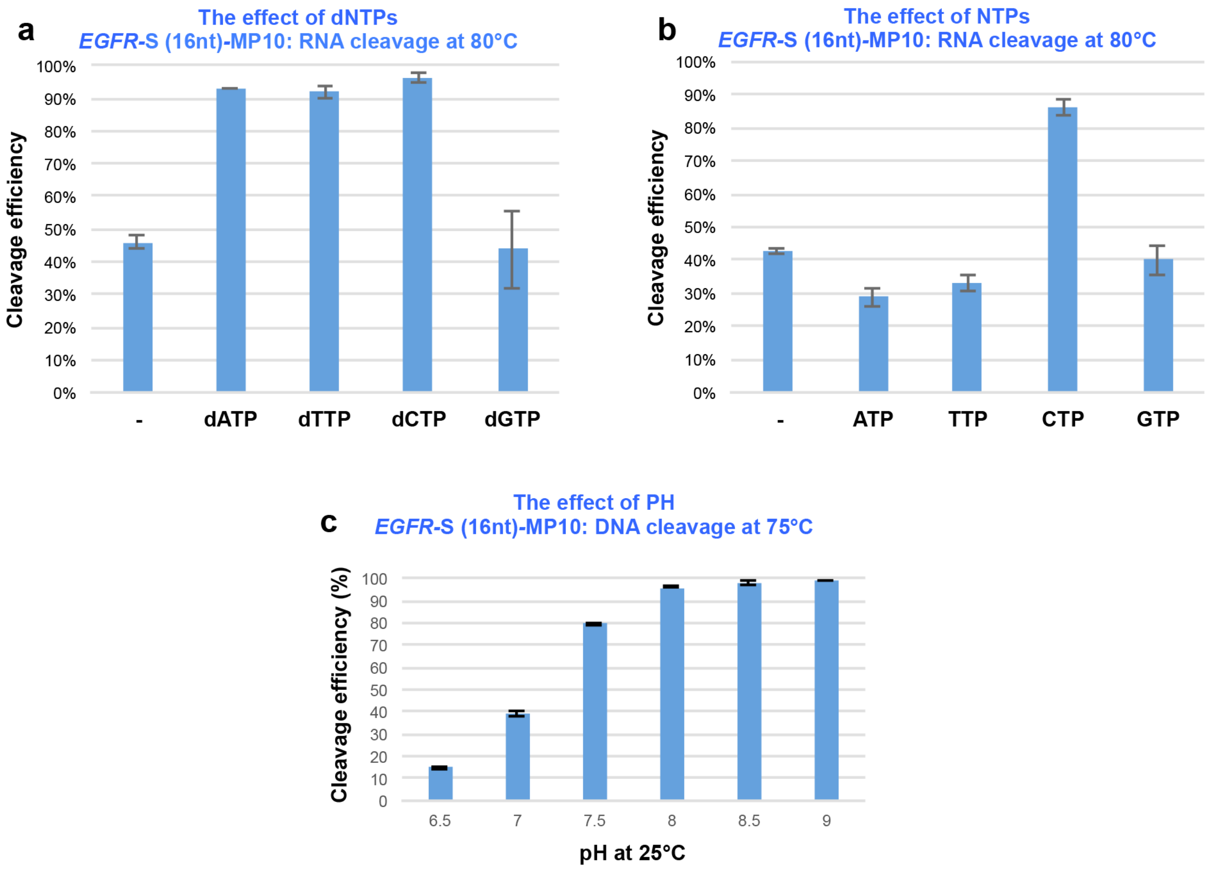


**Supplementary Figure 5:** The effects of **(a)** dNTPs and **(b)** NTPs on *EGFR* (L858R)-S RNA cleavage with guide *EGFR* (L858R)-S (16nt)-MP10 at 80 °C. **(c)** The effect of pH on *EGFR* (L858R)-S DNA cleavage with guide *EGFR* (L858R)-S (16nt)-MP10 at 75 °C. *Tt*Ago: guide: target =1: 0.2: 0.2. N=3.

### Supplementary Section 2: Single base pair mismatch discrimination

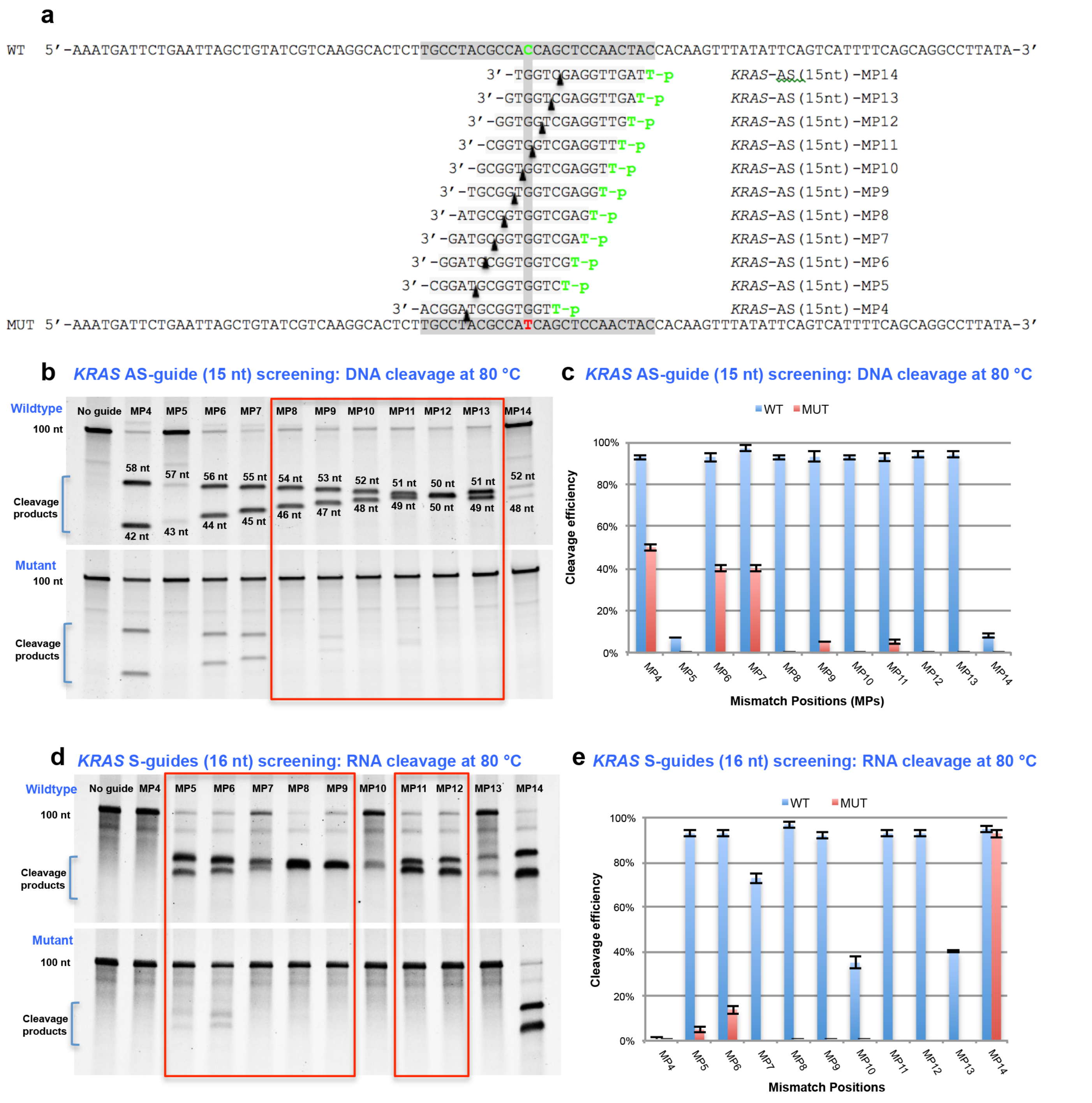


**Supplementary Figure 6: Cleavage efficiency of WT *KRAS* and *KRAS* G12D DNA and RNA as a function of MP.** (**a**) *KRAS* – antisense (AS) guide and AS target sequences. The various guides vary in the position of the pair mismatch between AS gDNA and AS *KRAS* G12D. (**b**) Electropherograms of cleaved AS WT *KRAS* and AS *KRAS* G12D DNA strands (80 $^{o}$C, 20 min). (**c**) Cleaving efficiencies of AS *KRAS* WT and AS *KRAS* G12D DNA as a function of MP. (**d**) Electropherograms of cleaved WT *KRAS* RNA and *KRAS* G12D RNA (80$^{o}$C, 20 min) (**e**) Cleaving efficiencies of WT *KRAS* RNA and *KRAS* G12D RNA as a function of MP. *Tt*Ago: guide: target =1: 0.2: 0.2. N=3.

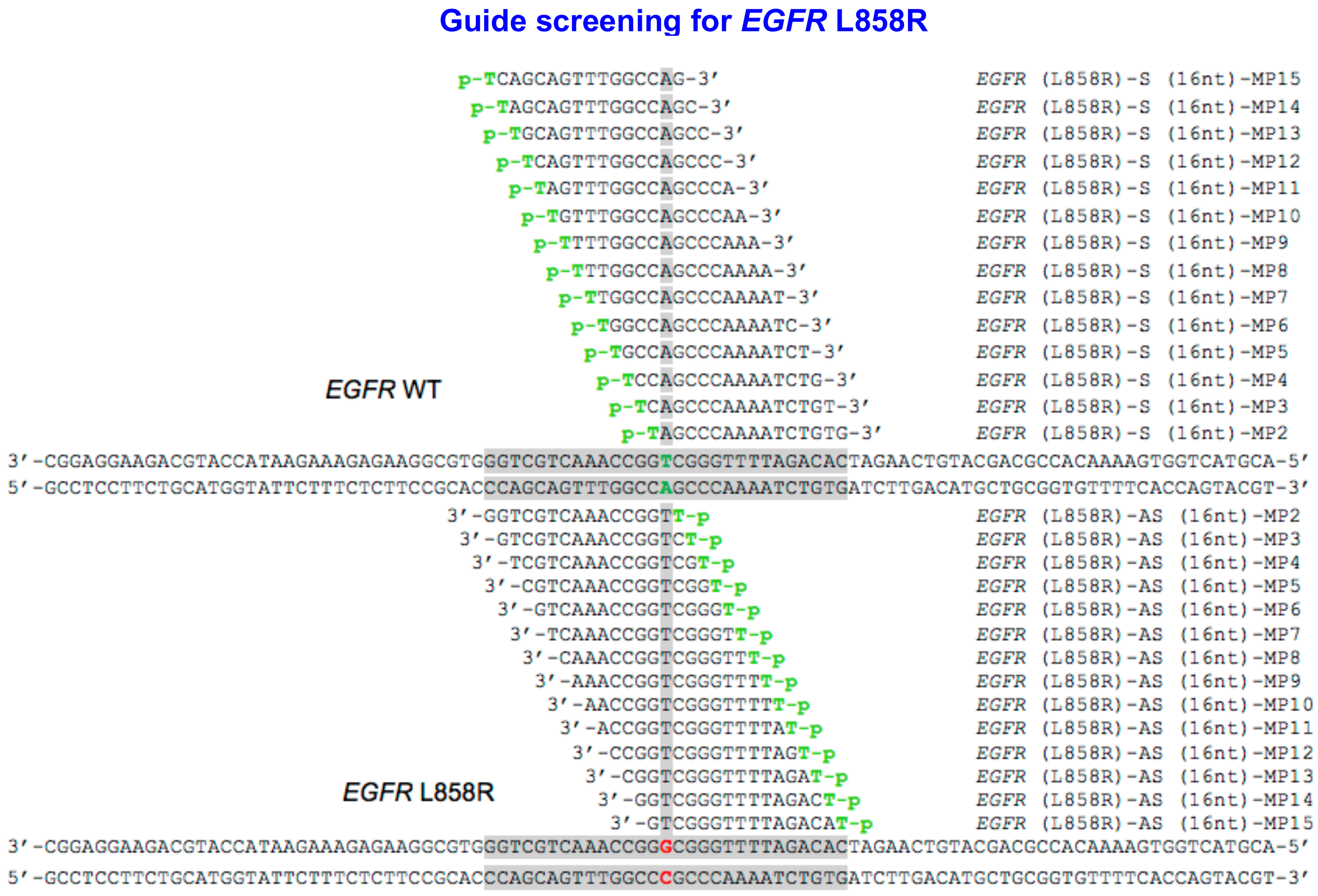


**Supplementary Figure 7: *EGFR* – guide and target sequences.** The various guides vary in the position of the pair mismatch between gDNA and *EGFR* L858R.

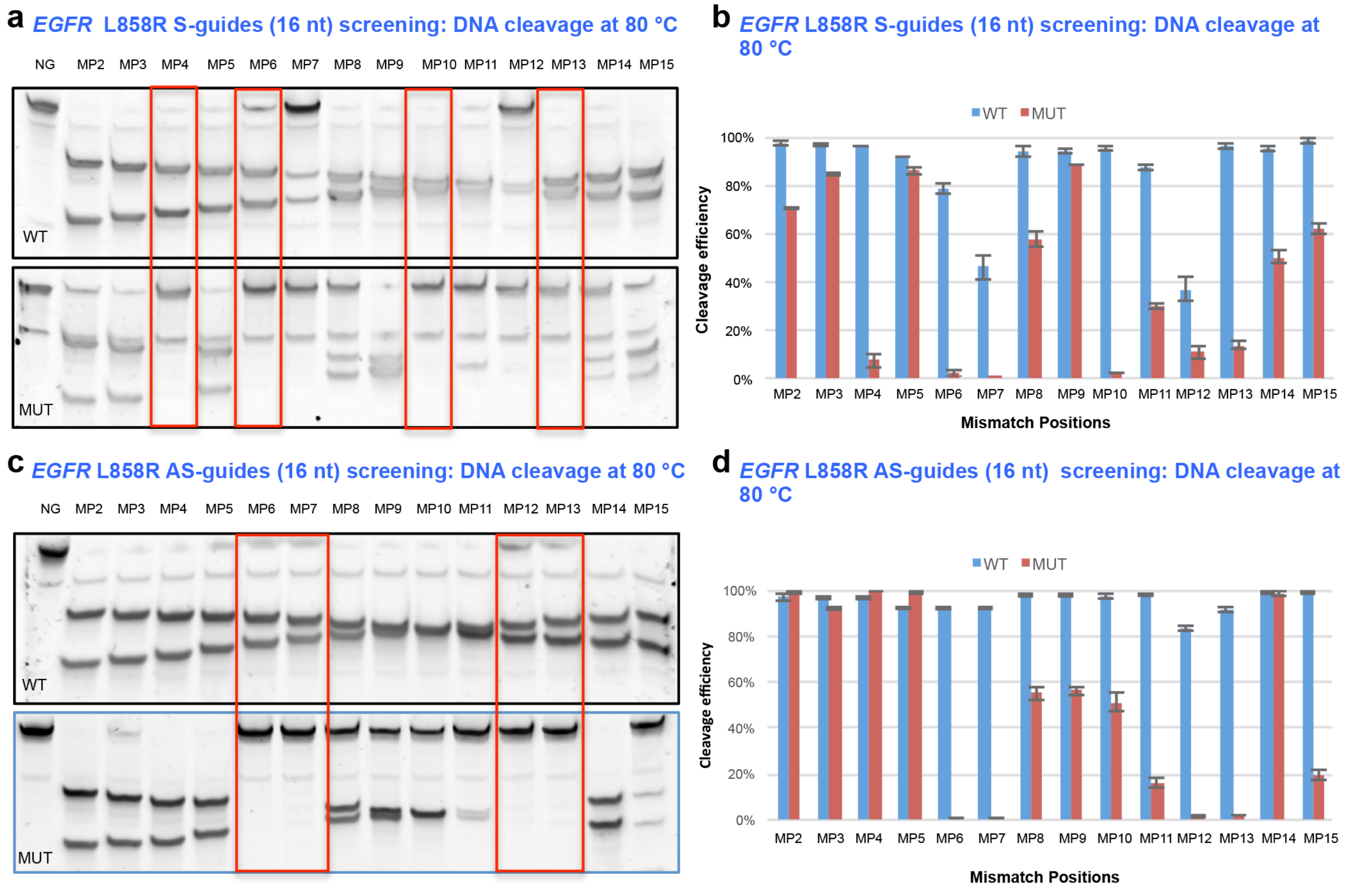


**Supplementary Figure 8: Cleavage efficiency as a function of the position of the mismatched pair between *EGFR* gDNA and *EGFR* L858R MA (DNA).** (**a**) Electropherograms of incubation products (80$^{o}$C, 20 min) of S WT *EFGR* and S *EFGR* L858R DNA strands. (**b**) Cleaving efficiency of S WT *EFGR* and S *EFGR* L858R DNA strands as a function of MP. (**c**) Electropherograms of incubation products (80 $^{o}$C, 20 min) of AS WT *EFGR* and AS *EFGR* L858R DNA strands. (**d**) Cleaving efficiency of AS WT *EFGR* and AS *EFGR* L858R DNA strands as a function of MP. *Tt*Ago: guide: target =1: 0.2: 0.2. N=3.

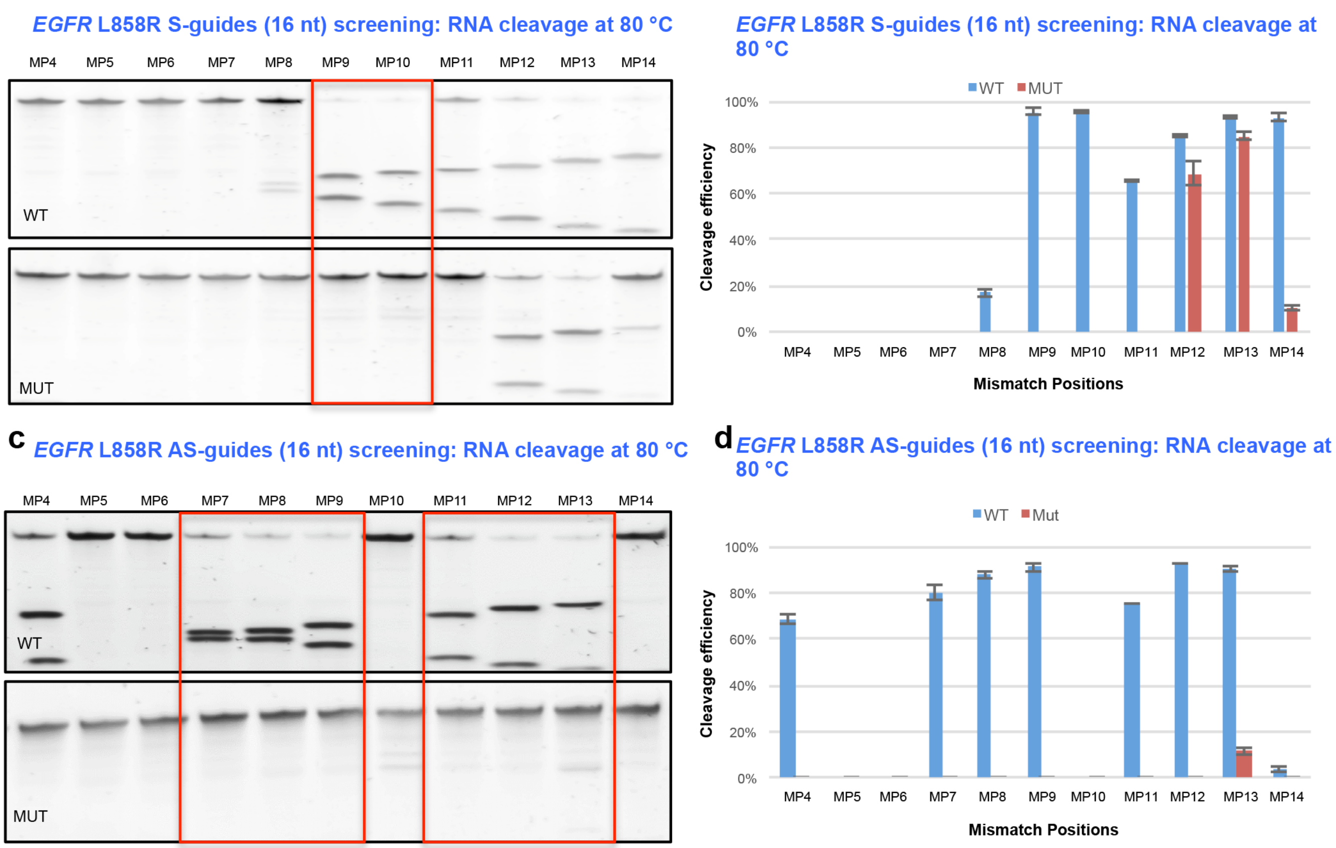


**Supplementary Figure 9: Cleavage efficiency as a function of the position of the mismatched pair between *EGFR* gDNA and *EGFR* L858R MA (RNA).** (**a**) Electropherograms of incubation products (80 °C, 20 min) of S WT *EFGR* and S *EFGR* L858R RNA strands. (**b**) Cleaving efficiency of S WT *EFGR* and S *EFGR* L858R RNA strands as a function of MP. (**c**) Electropherograms of incubation products (80 °C, 20 min) of AS WT *EFGR* and AS *EFGR* L858R RNA strands. (**d**) Cleaving efficiency of AS WT *EFGR* and AS *EFGR* L858R RNA strands as a function of MP. *Tt*Ago: guide: target =1: 0.2: 0.2. N=3.

### Guide screening for *EGFR* T790M

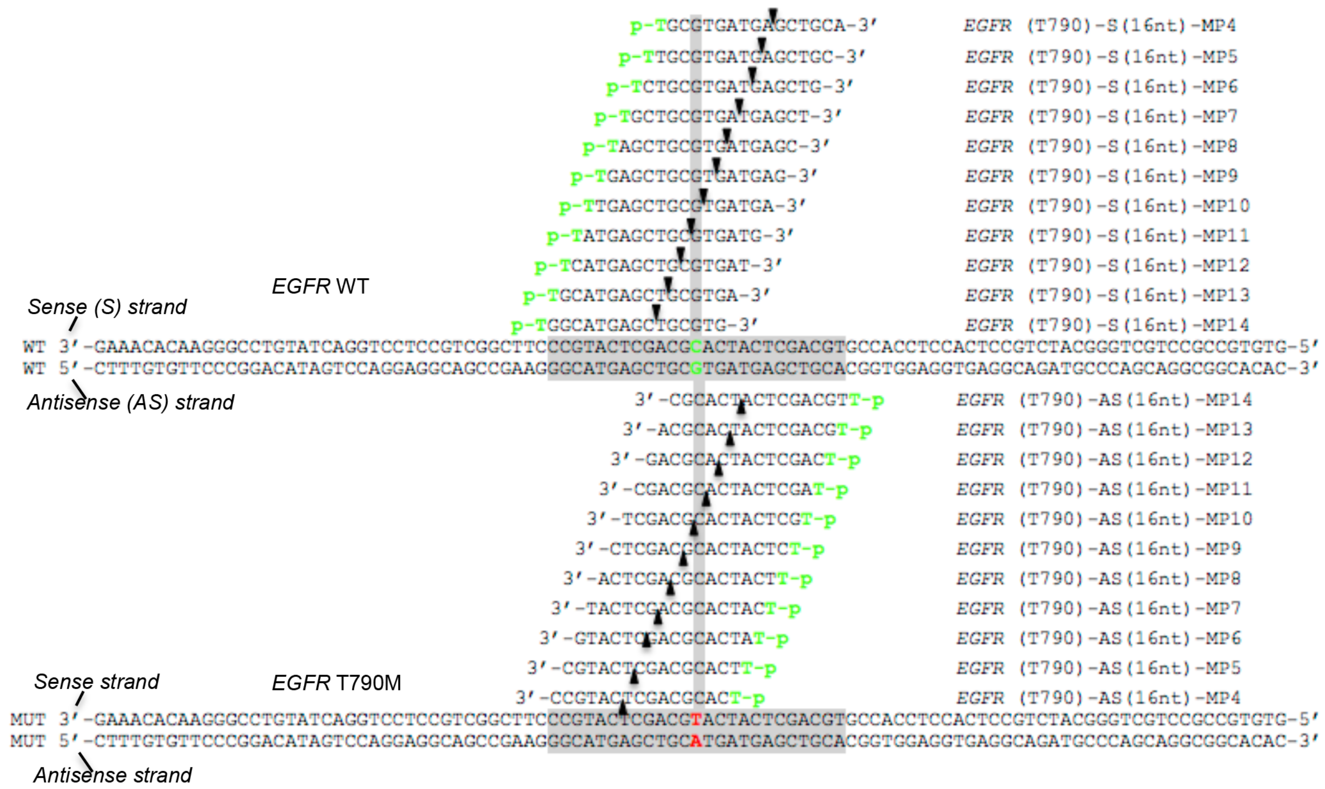


**Supplementary Figure 10: *EGFR* – guide and target sequences.** The various guides vary in the position of the pair mismatch between gDNA and *EGFR* T790M.

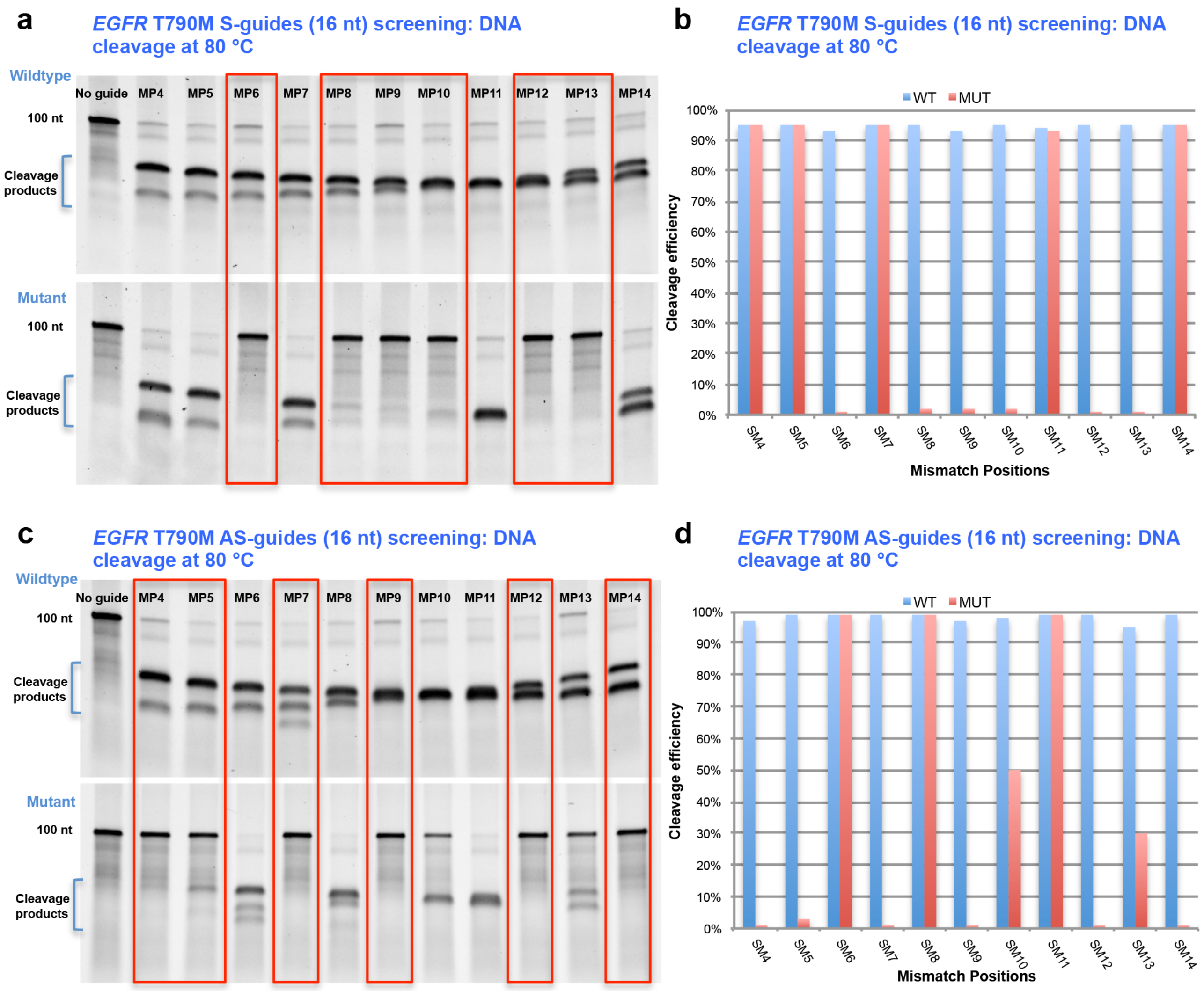


**Supplementary Figure 11: Cleavage efficiency as a function of the position of the mismatched pair between *EGFR* gDNA and *EGFR* T790M (DNA).** (**a**) Electropherograms of incubation products (80 $^{o}$C, 20 min) of S WT *EFGR* and S *EFGR* T790M DNA strands. (**b**) Cleaving efficiency of S WT *EFGR* and S *EFGR* T790M DNA strands as a function of MP. (**c**) Electropherograms of incubation products (80$^{o}$C, 20 min) of AS WT *EFGR* and AS *EFGR* T790M DNA strands. (**d**) Cleaving efficiency of AS WT *EFGR* and AS *EFGR* T790M DNA strands as a function of MP. *Tt*Ago: guide: target =1: 0.2: 0.2.

### Guide screening for *BRAF* V600E

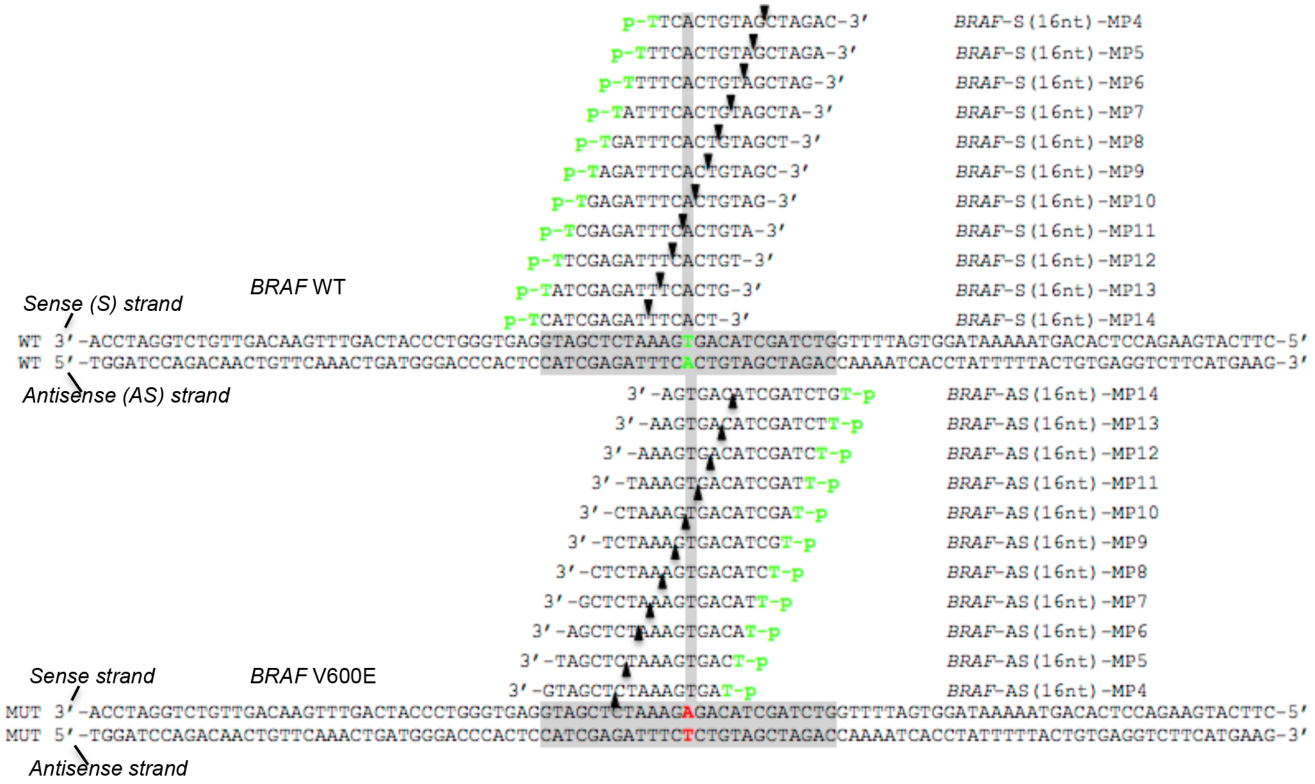


**Supplementary Figure 12: *BRAF* – guide and target sequences.** The various guides vary in the position of the pair mismatch between gDNA and *BRAF* V600E.

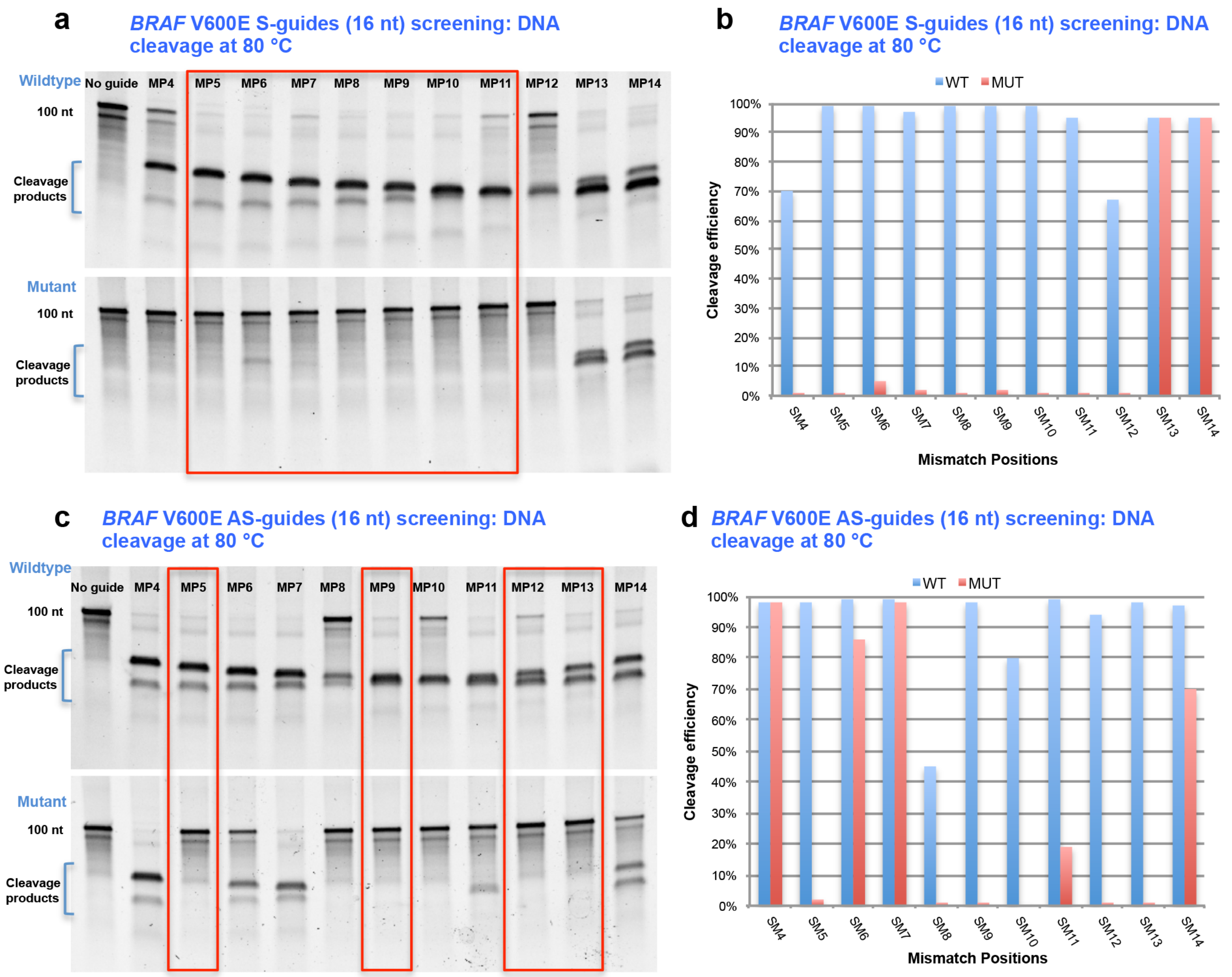


**Supplementary Figure 13: Cleavage efficiency as a function of the position of the mismatched pair between *BRAF* gDNA and *BRAF* V600E.** (**a**) Electropherograms of incubation products (80 °C, 20 min) of S WT *BRAF* and S *BRAF* V600E DNA strands. (**b**) Cleaving efficiency of S WT *BRAF* and S *BRAF* V600E DNA strands as a function of MP. (**c**) Electropherograms of incubation products (80 °C, 20 min) of AS WT *BRAF* and AS *BRAF* V600E DNA strands. (**d**) Cleaving efficiency of AS WT *BRAF* and AS *BRAF* V600E DNA strands as a function of MP. *Tt*Ago: guide: target =1: 0.2: 0.2.

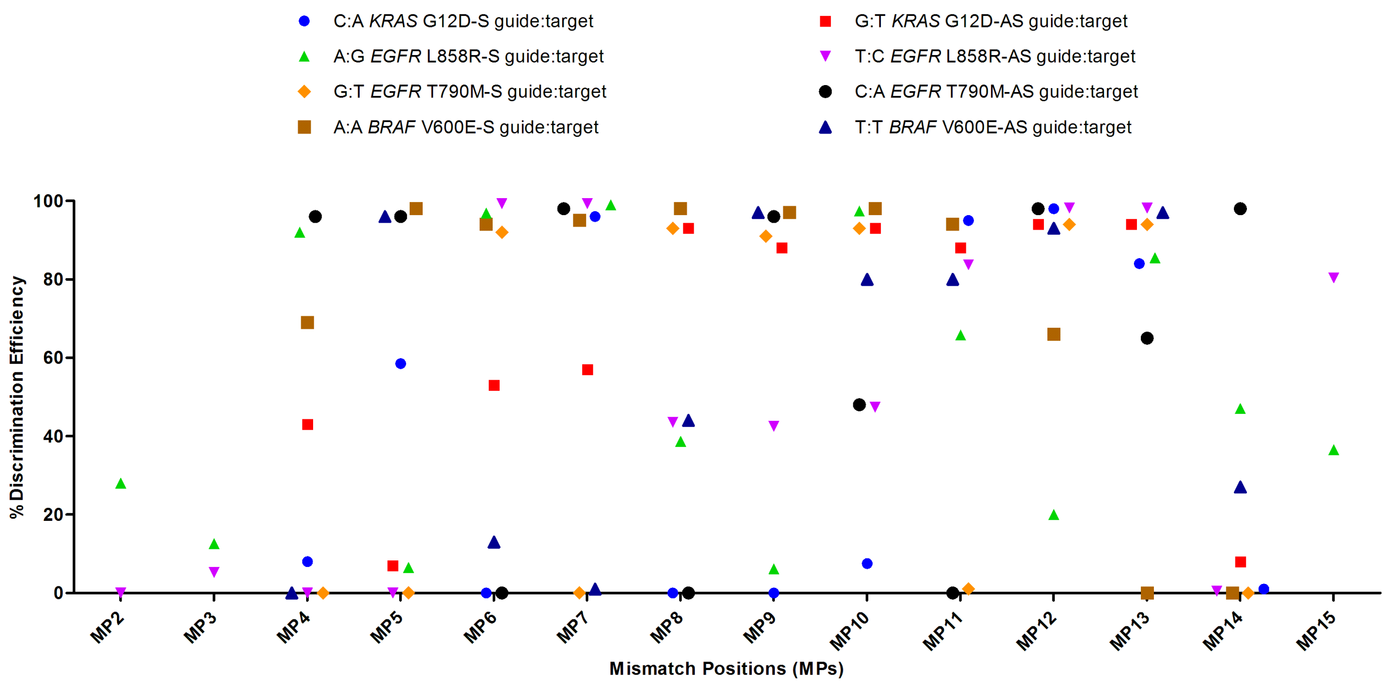


**Supplementary Figure 14: NAVIGATER discrimination efficiency depends sensitively on guide-off target mismatch pair's position.** Discrimination efficiency (the difference between cleaving efficiency of WA and MA) for S and AS *KRAS* G12D, *EGFR* L858R, *EGFR* T790M, and *BRAF* V600E DNA targets as a function of mismatched pair position.

**Supplementary Section 3: The effect of guide length and temperature on *Tt*Ago cleavage**

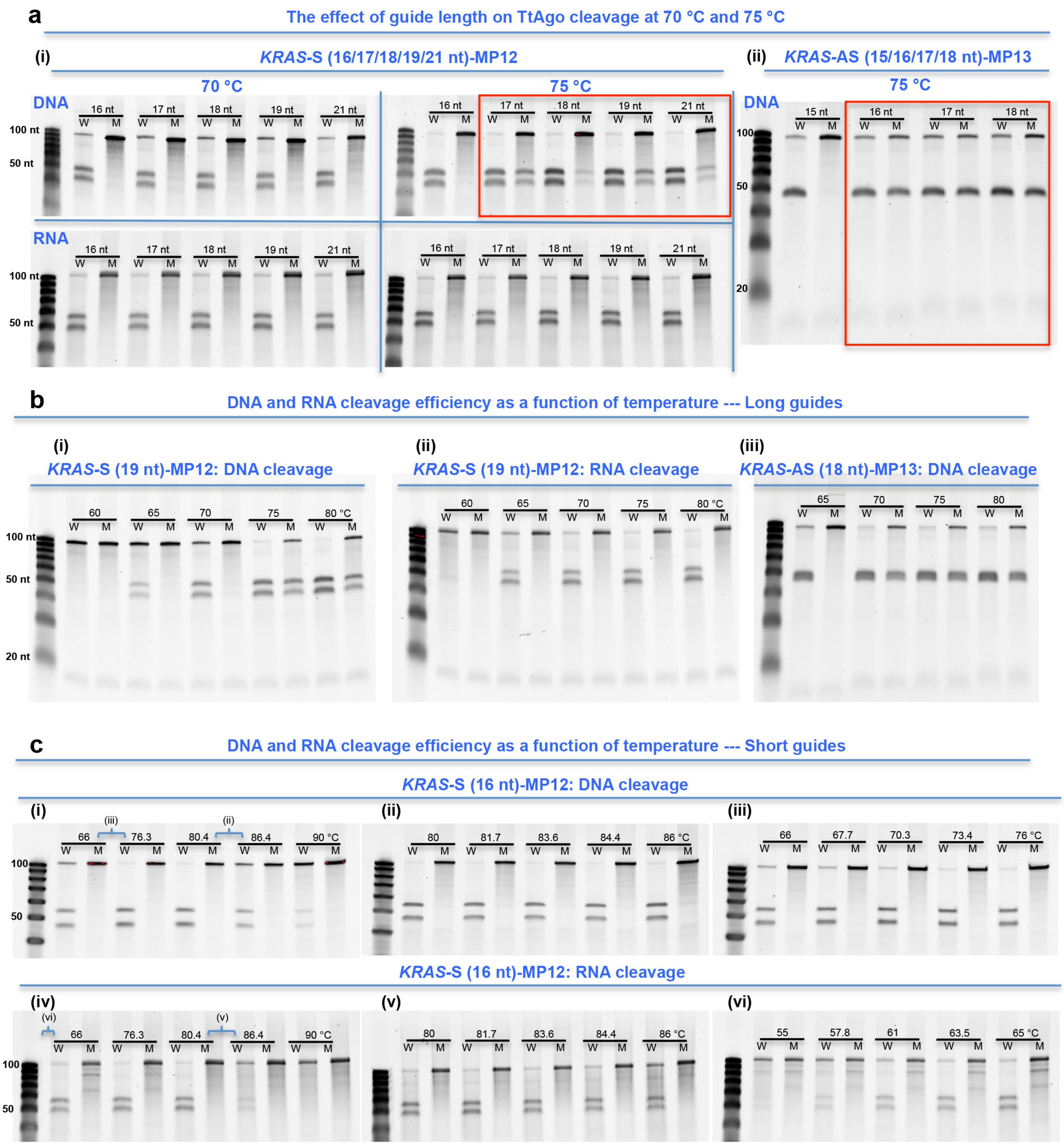


**Supplementary Figure 15: Electropherograms corresponding to Fig. 4. The combined effect of guide length and temperature on *Tt*Ago cleavage efficiency.** (a) The effect of guide length on *Tt*Ago cleavage at 70 and 75 °C. (i) Sense target strand *KRAS* DNA and RNA targets. (ii) Antisense target strand-*KRAS* DNA target. (b) Effect of temperature on cleavage efficiency using long (18/19 nt) guides with a guide/MA mismatch at positions g12 (S-*KRAS*) and g13 (AS *KRAS*). (c) *Tt*Ago cleavage with a short guide (16 nt) as a function of temperature. *Tt*Ago: guide: target =1: 0.2: 0.2. N=3.

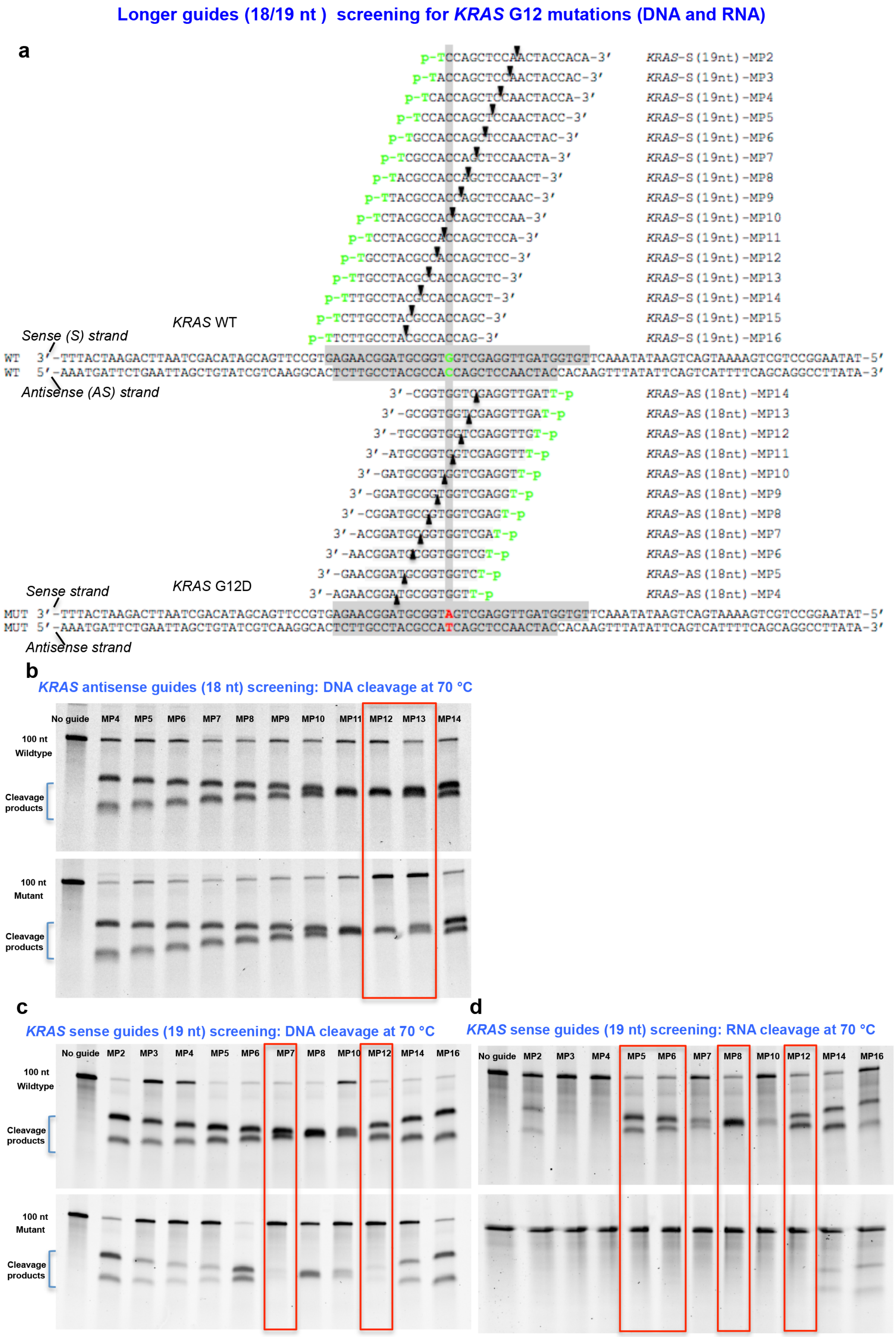


**Supplementary Figure 16: Cleavage efficiency of WT *KRAS* and *KRAS* G12D with long guides (18/19 nt) as a function of mismatched pair position.** (a) S (19 nt) and AS (18 nt)

guides' sequences. (b) AS KRAS 18 nt guide cleavage efficiency as a function of mismatched pair position at 70 $^{o}$C. (c) S *KRAS* 19 nt guide cleavage efficiency as a function of mismatched pair position at 70 $^{o}$C. *Tt*Ago: guide: target =1: 0.2: 0.2. N=3.

### Supplementary Section 4: dsDNA cleavage

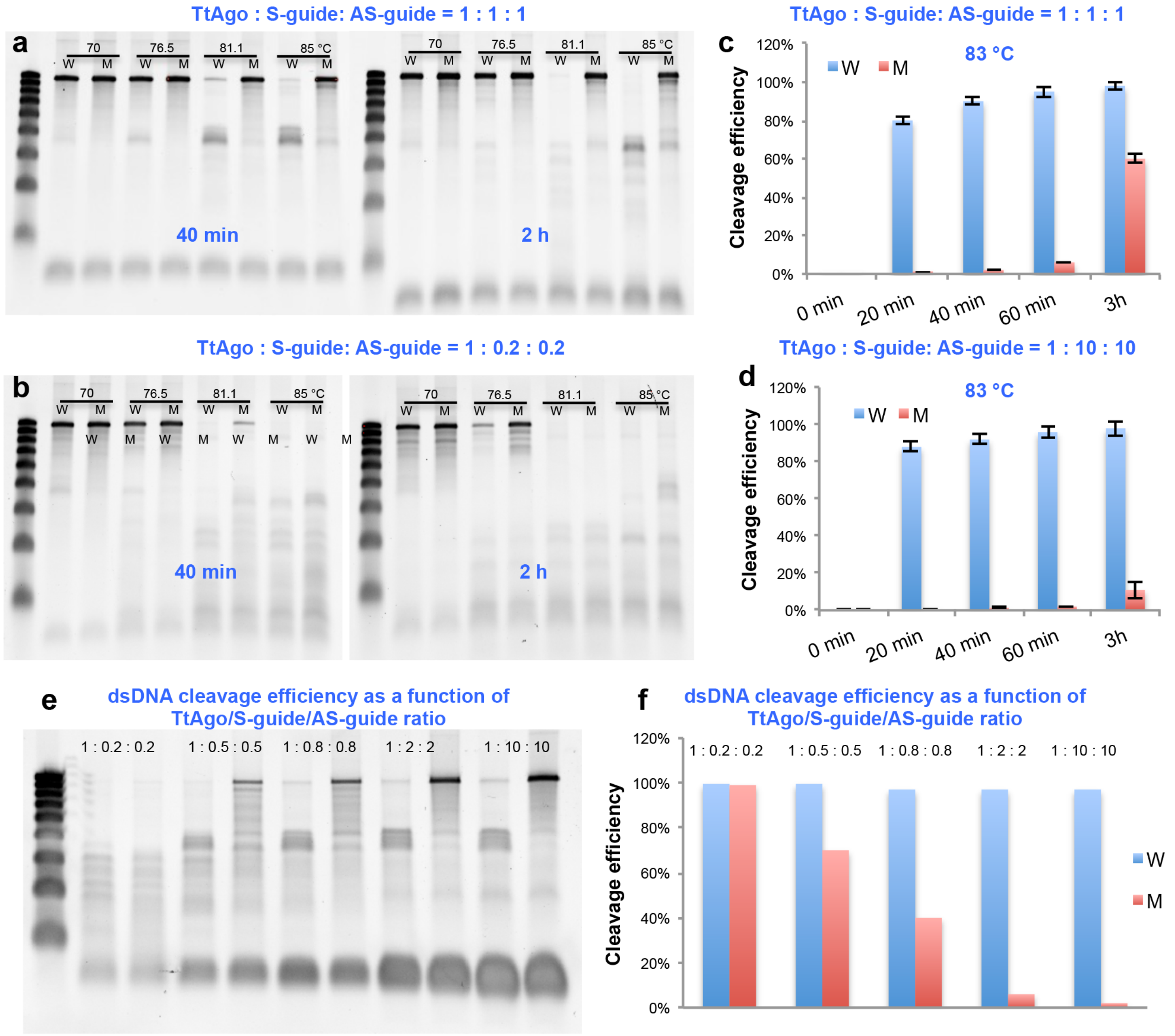


**Supplementary Figure 17: dsDNA cleavage efficiency as a function of temperature, guides concentrations, and incubation time. (a, b)** Electropherograms of ds*KRAS* (100 bp) cleaving assay products corresponding to **Figs. 5a** and **5b**. **(c)** Cleaving efficiency (corresponding to Fig. 5c**)** as a function of incubation time at 83 $^{o}$C. *Tt*Ago/S-guide/AS-guide ratio 1: 1: 1. **(d)** Cleaving efficiency (corresponding to Fig. 5d) as a function of incubation time at 83 $^{o}$C. *Tt*Ago/S-guide/AS-guide ratio 1: 10: 10. **(e)** Electropherograms of dsDNA cleavage as a function of *Tt*Ago/S-guide/AS-guide ratio. Incubation time 1h at 83 $^{o}$C. **(f)** Cleaving efficiency (corresponding to (e)) as a function of *Tt*Ago/S-guide/AS-guide ratio. All the experiments were carried out in Buffer 3 with *KRAS*-S (16nt)-MP12 and *KRAS*-AS (15nt)-MP13 guides.

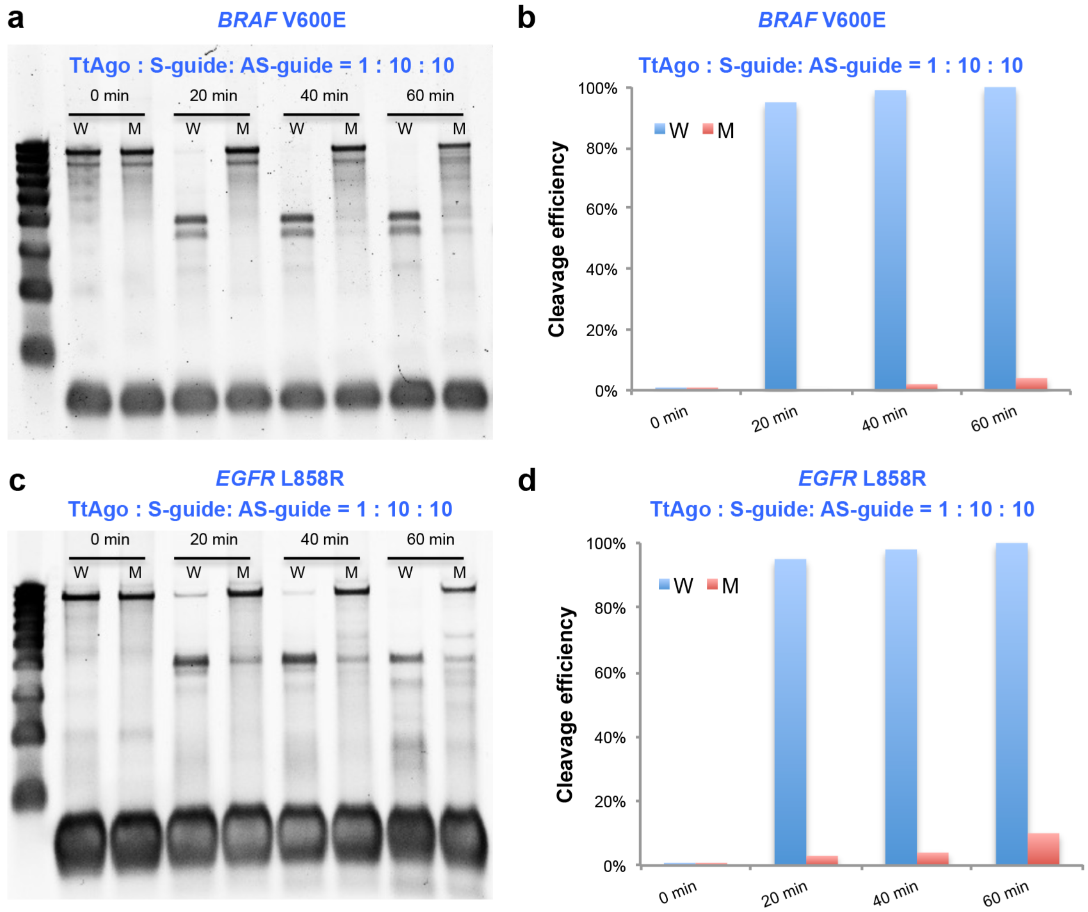


**Supplementary Figure 18: NAVIGATER's capability for enriching MAs harboring point mutations. (a)** Electropherograms of ds*BRAF* (100 bp) cleaving assay products as a function of incubation time. *BRAF*-S (16nt)-MP9 and *BRAF*-AS (16nt)-MP13 guides. **(b)** Cleaving efficiency of the assay in (a) as a function of incubation time. **(c)** Electropherograms of ds*EGFR* (100 bp) cleaving assay products as a function of incubation time. The experiment was carried out in buffer 3 with *EGFR*-S (16nt)-MP10 and *EGFR*-AS (15nt)-MP7 guides. **(d)** Cleaving efficiency of the assay in (c) as a function of incubation time. All experiments were carried out in Buffer 3 and 1 hour incubation time at 83$^{o}$C. *Tt*Ago/S-guide/AS-guide ratio 1: 10: 10.

### *EGFR* exon19 deletion mutations

**a**

| EGFR deletion (Ed) name | Frequency of EGFR exon19 deletion subtypes | Amino acid change | Base pair change | Length (bp) |
|---|---|---|---|---|
| **Ed1** | **45%** | p.E746_ A750del(1) | c.2235_2249del15 | 65 |
| **Ed2** | **19.6%** | p.E746_ A750del(2) | c.2236_2250del15 | 65 |
| **Ed3** | **8.4%** | p.L747_ P753>S | c.2240_2257del18 | 62 |
| **Ed4** | **4.3%** | p.L747_ T751del | c.2240_2254del15 | 65 |
| **Ed5** | **3.4%** | p.L747_ A750>P | c.2239_2250>cca | 71 |
| **Ed6** | **3.2%** | p.E746_ S752>V(2) | c.2237_2257>ttc | 62 |
| **Ed7** | **1.6%** | p.E746_ S752>V | c.2237_2255>t | 62 |
| **Ed8** | **1.4%** | p.L747_ S752del | c.2239_2256del18 | 62 |

**b**

```
EGFR deletion mutation-S-guide                3'-GTTCCTTAATTCTCTT-p
S WT (80bp):        5'-GGTGAGAAAGTTAAAATTCCCGTCGCTATCAAGGAATTAAGAGAAGCAACATCTCCGAAAGCCAACAAGGAAATCCTCGA-3'
S 2235_2249del15:   5'-GGTGAGAAAGTTAAAATTCCCGTCGCTATCAA---------------AACATCTCCGAAAGCCAACAAGGAAATCCTCGA-3'
S 2236_2250del15:   5'-GGTGAGAAAGTTAAAATTCCCGTCGCTATCAAG---------------ACATCTCCGAAAGCCAACAAGGAAATCCTCGA-3'
S 2240_2257del18:   5'-GGTGAGAAAGTTAAAATTCCCGTCGCTATCAAGGAAT------------------CGAAAGCCAACAAGGAAATCCTCGA-3'
S 2240_2254del15:   5'-GGTGAGAAAGTTAAAATTCCCGTCGCTATCAAGGAAT---------------CTCCGAAAGCCAACAAGGAAATCCTCGA-3'
S 2239_2250>cca:    5'-GGTGAGAAAGTTAAAATTCCCGTCGCTATCAAGGAA----CCA-----ACATCTCCGAAAGCCAACAAGGAAATCCTCGA-3'
S 2237_2257>ttc:    5'-GGTGAGAAAGTTAAAATTCCCGTCGCTATCAAGG------TTC------------CGAAAGCCAACAAGGAAATCCTCGA-3'
S 2237_2255>t:      5'-GGTGAGAAAGTTAAAATTCCCGTCGCTATCAAGG-------T-----------TCCGAAAGCCAACAAGGAAATCCTCGA-3'
S 2239_2256del18:   5'-GGTGAGAAAGTTAAAATTCCCGTCGCTATCAAGGAA------------------CCGAAAGCCAACAAGGAAATCCTCGA-3'

EGFR deletion mutation-AS-guide                        p-TAAGAGAAGCAACATC-3'
AS WT (80bp):       3'-CCACTCTTTCAATTTTAAGGGCAGCGATAGTTCCTTAATTCTCTTCGTTGTAGAGGCTTTCGGTTGTTCCTTTAGGAGCT-5'
AS 2235_2249del15:  3'-CCACTCTTTCAATTTTAAGGGCAGCGATAGTT---------------TTGTAGAGGCTTTCGGTTGTTCCTTTAGGAGCT-5'
AS 2236_2250del15:  3'-CCACTCTTTCAATTTTAAGGGCAGCGATAGTTC---------------TGTAGAGGCTTTCGGTTGTTCCTTTAGGAGCT-5'
AS 2240_2257del18:  3'-CCACTCTTTCAATTTTAAGGGCAGCGATAGTTCCTTA------------------GCTTTCGGTTGTTCCTTTAGGAGCT-5'
AS 2240_2254del15:  3'-CCACTCTTTCAATTTTAAGGGCAGCGATAGTTCCTTA---------------GAGGCTTTCGGTTGTTCCTTTAGGAGCT-5'
AS 2239_2250>cca:   3'-CCACTCTTTCAATTTTAAGGGCAGCGATAGTTCCTT----GGT-----TGTAGAGGCTTTCGGTTGTTCCTTTAGGAGCT-5'
AS 2237_2257>ttc:   3'-CCACTCTTTCAATTTTAAGGGCAGCGATAGTTCC------AAG------------GCTTTCGGTTGTTCCTTTAGGAGCT-5'
AS 2237_2255>t:     3'-CCACTCTTTCAATTTTAAGGGCAGCGATAGTTCC-------A-----------AGGCTTTCGGTTGTTCCTTTAGGAGCT-5'
AS 2239_2256del18:  3'-CCACTCTTTCAATTTTAAGGGCAGCGATAGTTCCTT------------------GGCTTTCGGTTGTTCCTTTAGGAGCT-5'
```

**c**

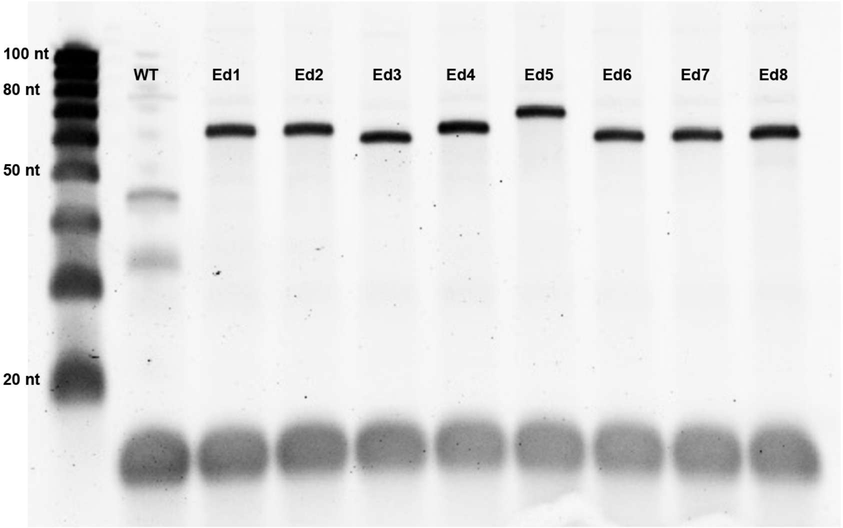


**Supplementary Figure 19: NAVIGATER's enriches MAs harboring deletion mutations. (a)** Common *EGFR* exon19 deletion mutations[1]. **(b)** Target sequences and guides for enriching MAs containing *EGFR* exon19 deletion mutations. **(c)** Electropherograms of cleaving assay products of WT ds*EGFR* (80 bp) and ds*EGFR* exon19 deletion mutants. Incubation time 1 hour at 83$^{o}$C. *Tt*Ago/S-guide/AS-guide ratio 1: 10: 10. Synthetic dsDNAs harboring common *EGFR* exon19 deletion mutations were used.

### Supplementary Section 4: NAVIGATER improves the sensitivity of downstream rare allele detection

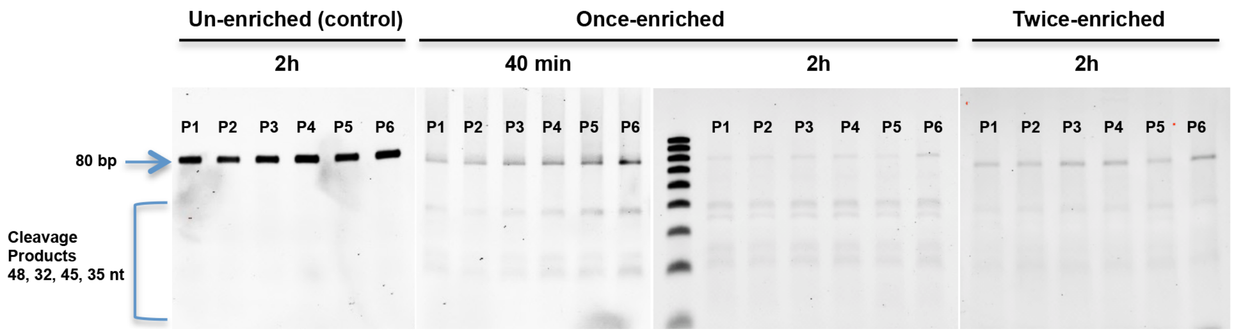


**Supplementary Figure 20: Electropherograms of six pancreatic cancer patient's samples (Table 2) without enrichment (control), once enriched for 40 min and 2h and twice-enriched.**

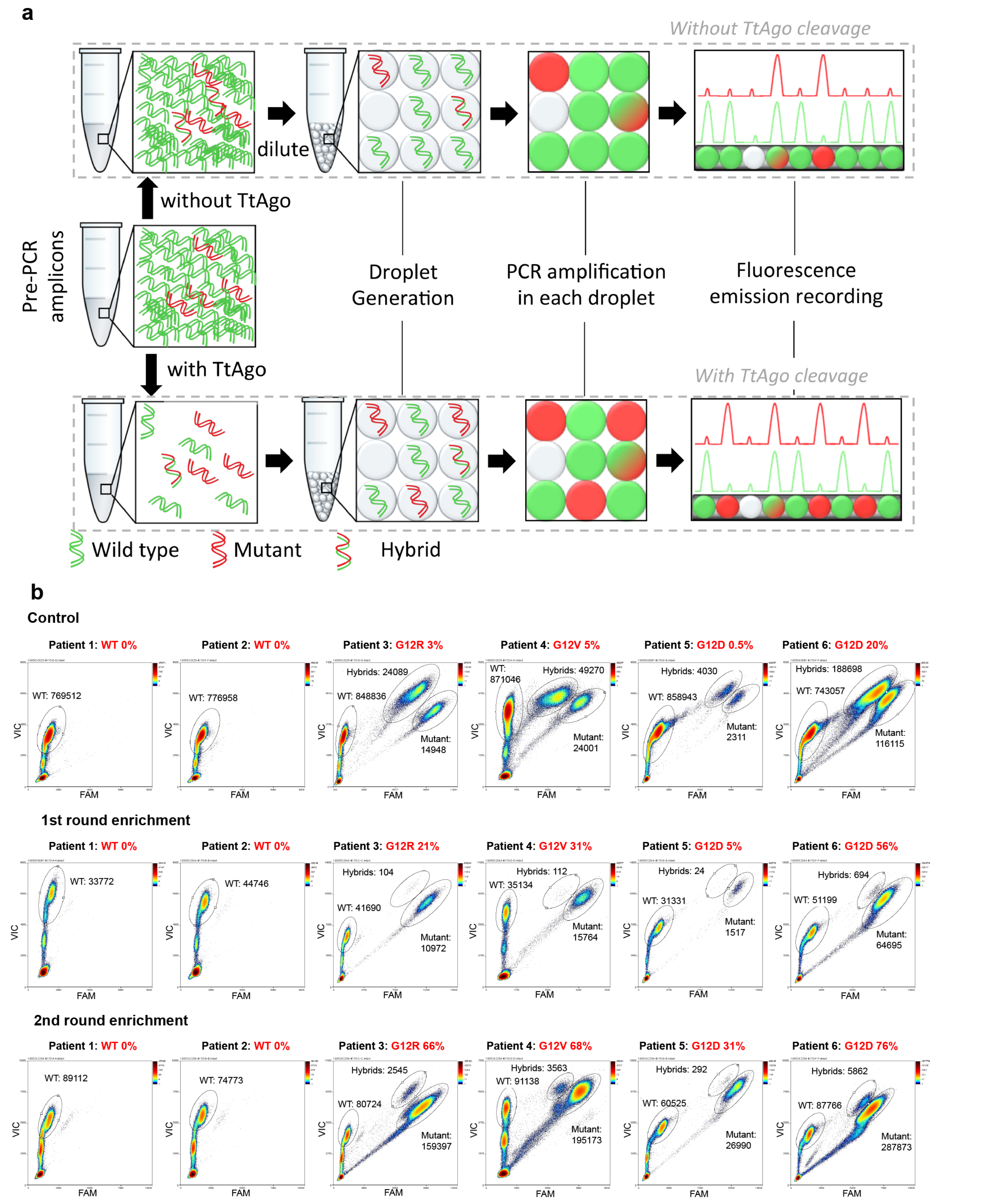


**Supplementary Figure 21: Six pancreatic cancer patient's samples processed with NAVIGATER and ddPCR.** (a) Principle of operation of NAVIGATER combined with ddPCR. (b)

ddPCR results of non-enriched, once-enriched, and twice-enriched samples. There are three signaling droplet populations for *KRAS*: WT, mutant, and WT-mutant hybrid. 2~7 millions of droplets are produced for each sample.

**Supplementary Section 5: Multiplexed enrichment and detection with XNA-PCR**

**Results**

In the absence of *Tt*Ago treatment (control), the electropherograms of single-plex and triplex PCR products feature five bands, one at ~120 bp (predominantly WT *KRAS* G12D amplicon), one at ~100 bp (predominantly, WT *EGFR* sequence susceptible to L858R mutation, and WT *EGFR* sequence susceptible to ΔE746 - A750 mutation), one at ~75 bp (internal control amplicon), one at ~30 bp (Taqman probe), and one at 16 nt (gDNA) (**Supplementary Fig. 22a**).

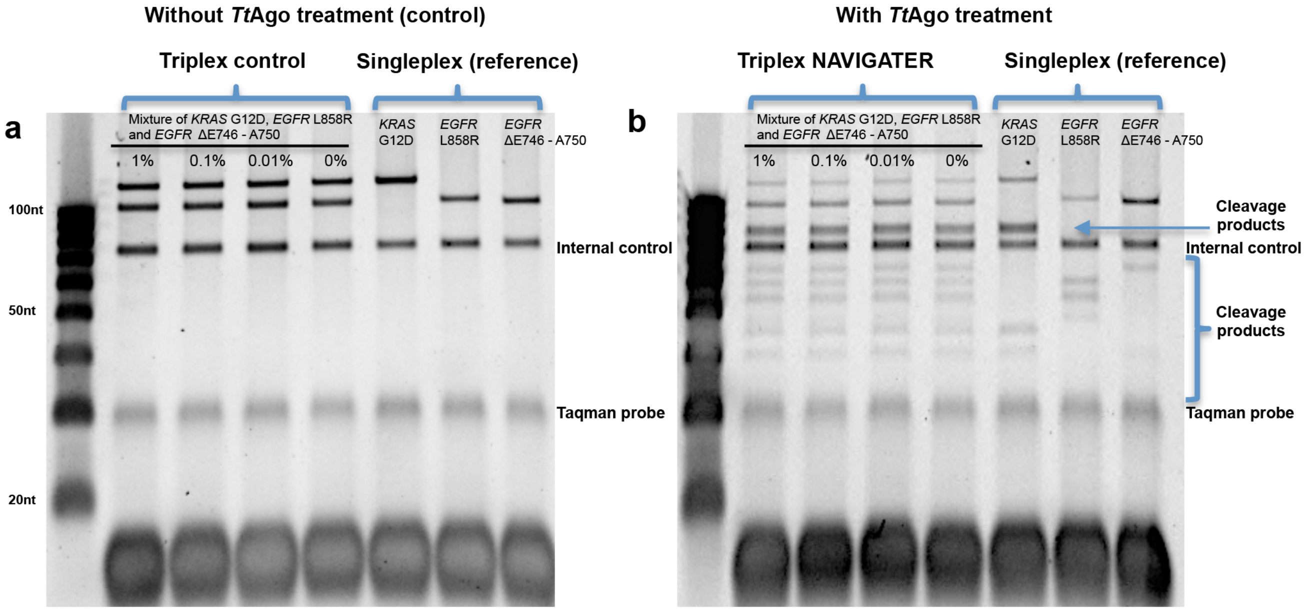


**Supplementary Figure 22: Multiplexed enrichment of *KRAS* G12D, *EGFR* L858R, and *EGFR* ΔE746 - A750.** Electropherograms of triplex PCR products without (**a**) and with (**b**) NAVIGATER pre-enrichment.

After 1 hour incubation, with NAVIGATER, the bands at ~120 bp and ~100 bp faded, indicating a reduction of WT *KRAS* and WT *EGFR* alleles, while there is no change in the band intensities of the internal control (**Supplementary Fig. 22b**). The one-to-one correspondence between the electrophrograms of the triplex assays and singleplex assays indicates that there is no interference among reagents, and that NAVIGATER provides specific enrichment and is amenable to multiplexing (**Supplementary Fig. 22b**).

Furthermore, we tested NAVIGATER's ability to improve detection sensitivity of the mutation detection assay XNA-PCR[5]. We processed samples of 60 ng cfDNA that contain various fractions of *KRAS* G12D, *EGFR* ΔE746 - A750, and *EGFR* L858R. In the absence of pre-enrichment, XNA-PCR can detect 0.1% *KRAS* G12D, 0.1% *EGFR* ΔE746 - A750, and 1% *EGFR* L858R but fails to detect 0.01% *KRAS* G12D, 0.01% *EGFR* ΔE746 - A750, and 0.1% *EGFR* L858R (**Supplementary Figs. 23a, 23c, and 23e**). After NAVIGATER pre-enrichment, XNA-PCR detects readily 0.01% *KRAS* G12D, 0.01% *EGFR* ΔE746 - A750, and 0.1% *EGFR* L858R (**Supplementary Figs. 23b, 23d, and 23f**). NAVIGATER pre-enrichment can improve XNA-PCR sensitivity by at least an order of magnitude.

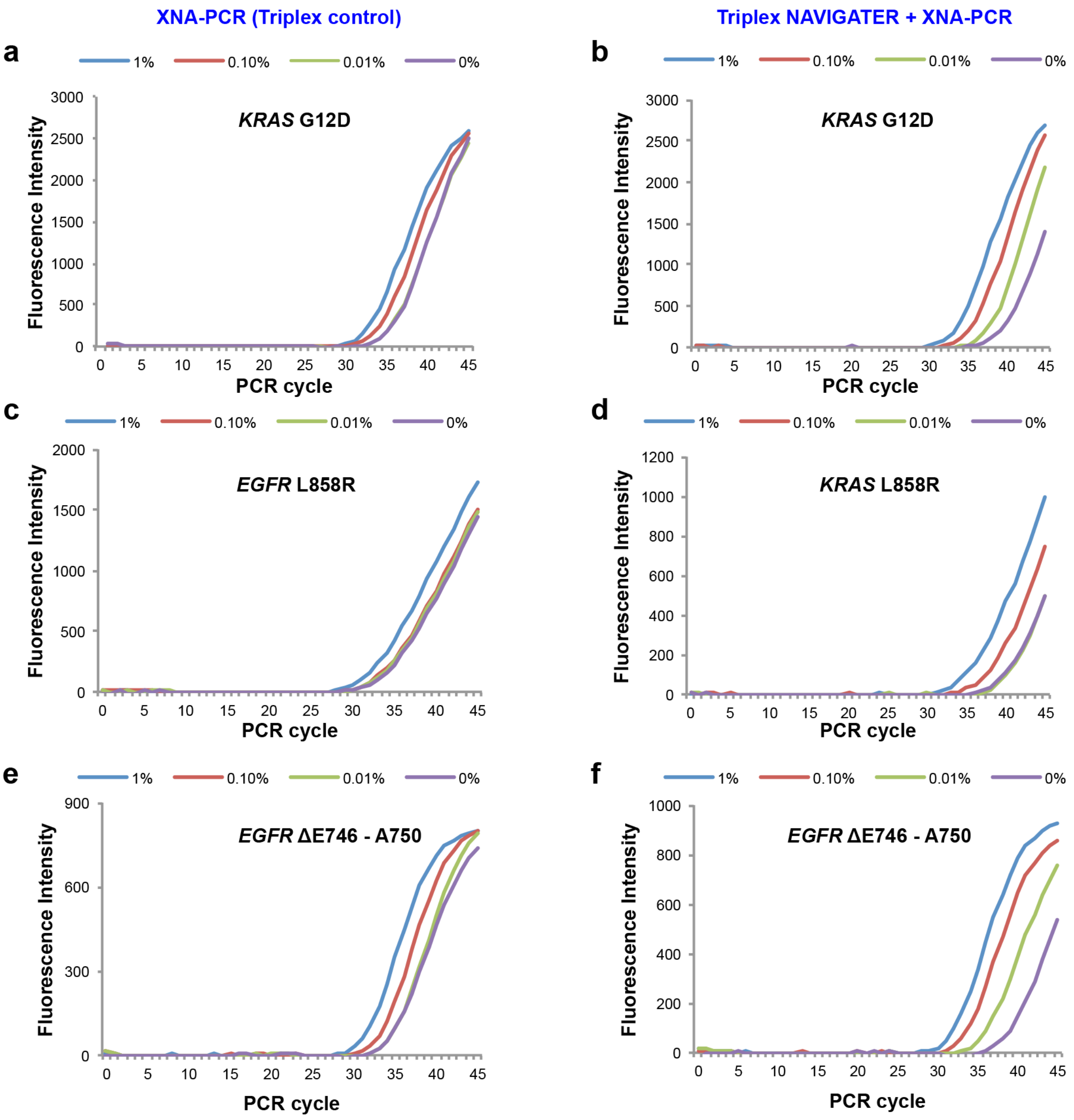


**Supplementary Figure 23: Multiplexed enrichment of *KRAS* G12D, *EGFR* L858R, and *EGFR* ΔE746 - A750.** XNA-PCR amplification curves of samples of various fractions of *KRAS* G12D, *EGFR* L858R and *EGFR* ΔE746 - A750 in the absence (a, c, e) and presence (b, d, f) of NAVIGATER pre-treatment. Triplex NAVIGATER increases the sensitivity of XNA-PCR by at least an order of magnitude. All the controls were pre-processed with NAVIGATER in the absence of *Tt*Ago.

### Supplementary Section 6: TtAgo Purification

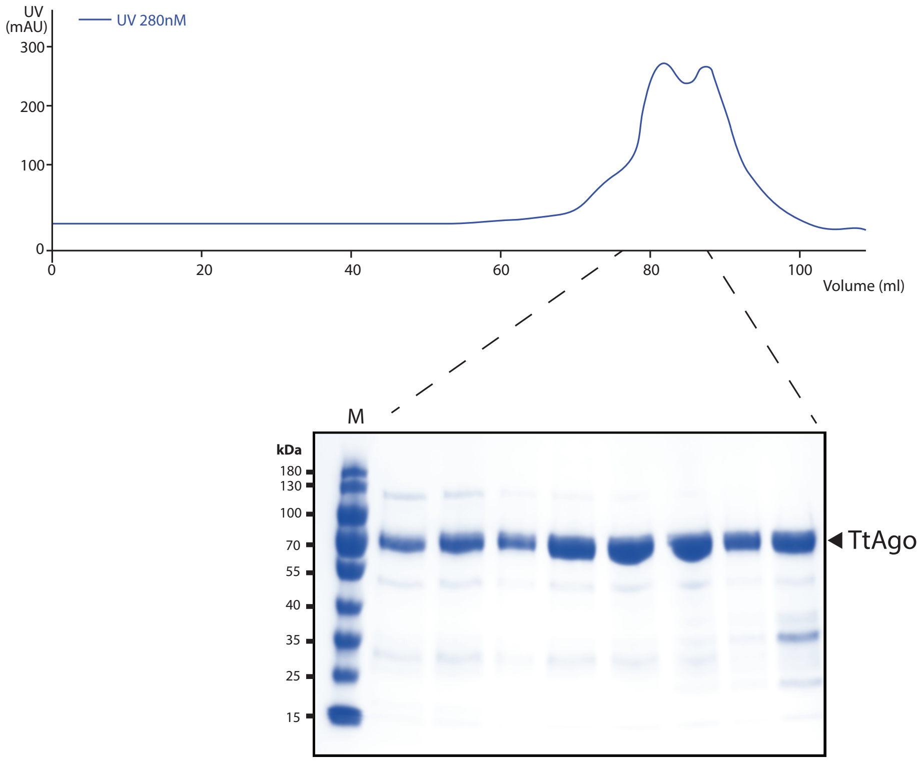


**Supplementary Figure 24: Chromatopherogram of purified *Tt*Ago.**

**Supplementary Table 1:** The sequences and concentrations of *KRAS* primers, PNA clamp oligo, and Taqman probes used in downstream mutation analysis.

| Name | Sequences (5'---3') | Concentration | Amplicons length and melting temperature |
|---|---|---|---|
| 80 bp-PCR-FW[2] | AGGCCTGCTGAAAATGACTGAATAT | | 80 bp, 79.5°C |
| 80 bp-PCR-RV[2] | GCTGTATCGTCAAGGCACTCTT | | |
| G12-WT-VIC probe[2] | TTGGAGCTGGTGGCGT | 100 nM | |
| G12D-FAM probe[2] | TGGAGCTGATGGCGT | 100 nM | |
| G12R-FAM probe[2] | TTGGAGCTCGTGGCGT | 100 nM | |
| G12V-FAM probe[2] | ACGCCAACAGCTC | 100 nM | |
| 295 bp-PCR-FW[3] | AAGGTACTGGTGGAGTATTTG | 100 nM | 295 bp, 82.6°C |
| 295 bp-PCR-RV[3] | GTACTCATGAAAATGGTCAGAG | 100 nM | |
| SMAP-2-OP1[4] | TATTATAAGGCCTGCTG | 0.4 μM | 123 bp |
| SMAP-2-OP2[4] | TTGGATCATATTCGTCC | 0.4 μM | |
| SMAP-2-FP[4] | ACCTTCTACCCTCAGAAGGTATAAACTTGTG GTAGTTGGAGC | 3.0 μM | |
| SMAP-2-BP[4] | GCAAGAGTGCCTTGA | 1.5 μM | |
| SMAP-2-TP[4] | TGGCGTAGGCATGATTCTGAATTAGCTGTAT | 3.0 μM | |
| PNA clamp[4] | CCTACGCCACCAGCTCC | 0.7 μM | |
